# Microfluidic low-input profiling reveals lncRNA roles in disease

**DOI:** 10.64898/2026.03.19.712992

**Authors:** Jenna A. Catalano, Yuan-Pang Hsieh, Zhengzhi Liu, Gaoshan Li, J. Javier Meana, Javier González-Maeso, Zhen Bouman Chen, Chang Lu

## Abstract

Long noncoding RNAs (lncRNAs) regulate gene expression through binding to DNA, various RNAs, and proteins, playing potentially important but poorly understood roles in diseases. Existing approaches for profiling lncRNA–chromatin interactions at the genome scale require large quantities of input material (e.g., 100 million cells per assay). Applying these technologies to tissue samples has been challenging especially when examination of a specific cell type is desired. Here we demonstrate a low-input microfluidic technology based on Chromatin Isolation by RNA Purification (ChIRP) for mapping lncRNA–chromatin interactions using as few as 50,000 cells. We validate our technology, muChIRP-seq, on two lncRNAs of different sizes (*GOMAFU* and *TERC*) in human and mouse cell lines and in brain tissues. Furthermore, we profile neuronal nuclei from postmortem human brain tissues of schizophrenia and control subjects. Our profiling data reveal distinct roles and levels of involvement for the two lncRNAs in contribution to schizophrenia. Our multimodal integrative analysis suggests coordination between lncRNA binding and other epigenomic mechanisms such as histone modifications in schizophrenia pathogenesis. Our technology enables lncRNA studies in tissue samples and in a cell-type-specific fashion, unlocking new opportunities to screen and understand lncRNA involvement in diseases.

## Introduction

Only 1.5% of the human genome encodes proteins, while the remainder is transcribed into various noncoding RNA^1–4^. Long noncoding RNAs (lncRNAs) are transcripts longer than 200 bp that regulate gene expression through their interactions with DNA, other RNAs, and proteins during diseases and development^1–6^. However, how lncRNAs carry out their regulatory functions by interacting with the genome remains largely unknown.

Two major approaches have been taken to study lncRNA-chromatin interactions in the past. Affinity-based approaches like ChIRP-seq^7–10^, CHART-seq^11^, and RAP-seq^12,13^ focus on studying the binding profile of a single lncRNA by targeting its sequence. In these methods, oligonucleotide probes are designed to capture lncRNA-bound chromatin in a lncRNA sequence-specific manner. Alternatively, proximity-based methods like MARGI-seq^14–16^, GRID-seq^17,18^, CHAR-seq^19,20^, RADICL-seq^21^, and MUSIC-seq^22^ establish RNA and DNA interactions for an “all-to-all” mapping of the chromatin landscape. Affinity-based methods allow for mapping of a single lncRNA of known sequence, whereas proximity-based methods allow mapping of novel lncRNAs and mutations. Proximity-based methods typically require more sequencing resource, and their data tends to be biased toward highly abundant lncRNA.

Both types of approaches are limited by requiring a large input of starting material. Most of these methods require tens to a hundred million cells for each assay, with the exception of iMARGI-seq^15,16^, RADICL-seq^21^, and MUSIC-seq^22^, which require millions of input cells. Due to the input limitation, most previous studies involving lncRNA profiling were conducted using cell lines^7,11,13,14,16,17,19,21^. Experimentation on tissue samples with direct biomedical relevance has been scarce^22,23^. Furthermore, similar to other epigenetic regulatory mechanisms, lncRNAs are known to function in a cell-type and disease-specific manner and their binding profiles vary amongst individuals^1–6,24–26^. Thus low-input methods that permit profiling genome-wide lncRNA-chromatin interactions with cell-type specificity are critical for future large-scale functional studies of lncRNAs in tissue samples, in the spirit of encyclopedia of DNA elements (ENCODE) effort^27^, and for investigating lncRNA involvement in diseases.

Here we demonstrate a microfluidic technology referred to as muChIRP-seq for low-input affinity-based approach to studying lncRNA binding using as few as 50,000 cells. We first validated our approach against published ChIRP-seq data for a lncRNA *TERC*^7^. We then collected data on *Gomafu* (*Miat*), a lncRNA linked with cardiac health^28,29^, cancer^30,31^, and mental illness^32–35^ whose binding had not been profiled at the genome scale prior to this work. Furthermore, we applied our technology to study lncRNA binding for both *GOMAFU* and *TERC* in NeuN+ postmortem human prefrontal cortex nuclei from schizophrenia and control patients. Our data revealed distinct levels of involvement for the two lncRNAs in schizophrenia. MuChIRP-seq data were integrated with RNA-seq, H3K4me3, and H3K27ac data^36^ to understand the regulatory roles of these lncRNAs. Our technology offers a scalable approach that enables large-scale studies of lncRNA binding using tissue samples with direct disease relevance.

## Results

### MuChIRP-seq workflow

MuChIRP utilizes a microfluidic device fabricated using multilayer soft lithography^37^ to improve ChIRP^7,8^ capture efficiency (**Figure 1a** and **1b**, **Supplementary Figure 1**). The device has a single microfluidic chamber (∼5.7 μL) with a two-layer pneumatic sieve valve at the outlet^38,39^. Several steps are involved in the muChIRP process. First, cells are fixed using glutaraldehyde to immobilize lncRNA binding to the genome before sonication is conducted to break the genome into 100-500 bp fragments that contain lncRNA-chromatin complexes **(Supplementary Figure 2)**. Second, streptavidin C-1 magnetic beads are coated with biotinylated tiling oligonucleotide probes that target a specific lncRNA and are separated into “even” and “odd” probe pools^7,8^. Third, the chromatin fragments undergo a two-step hybridization with probe-coated beads in the microfluidic device. In the oscillatory hybridization step, magnetic beads are held in place by a magnet while chromatin fragments are flowed back and forth through the beads for 4 h, driven by automated pressure pulses at the two ends of the chamber. The beads are then packed against the closed sieve valve and chromatin is slowly flowed through the packed bed of beads to enable collection of additional lncRNA-associated chromatin. Fourth, an oscillatory washing step removes non-specifically bound chromatin from the beads. Finally, beads bearing lncRNA-associated chromatin are collected for off-chip processing. Six muChIRP-seq devices are fixed to a single glass slide for easy parallelization (**Figure 1b**).

**Figure 1.**
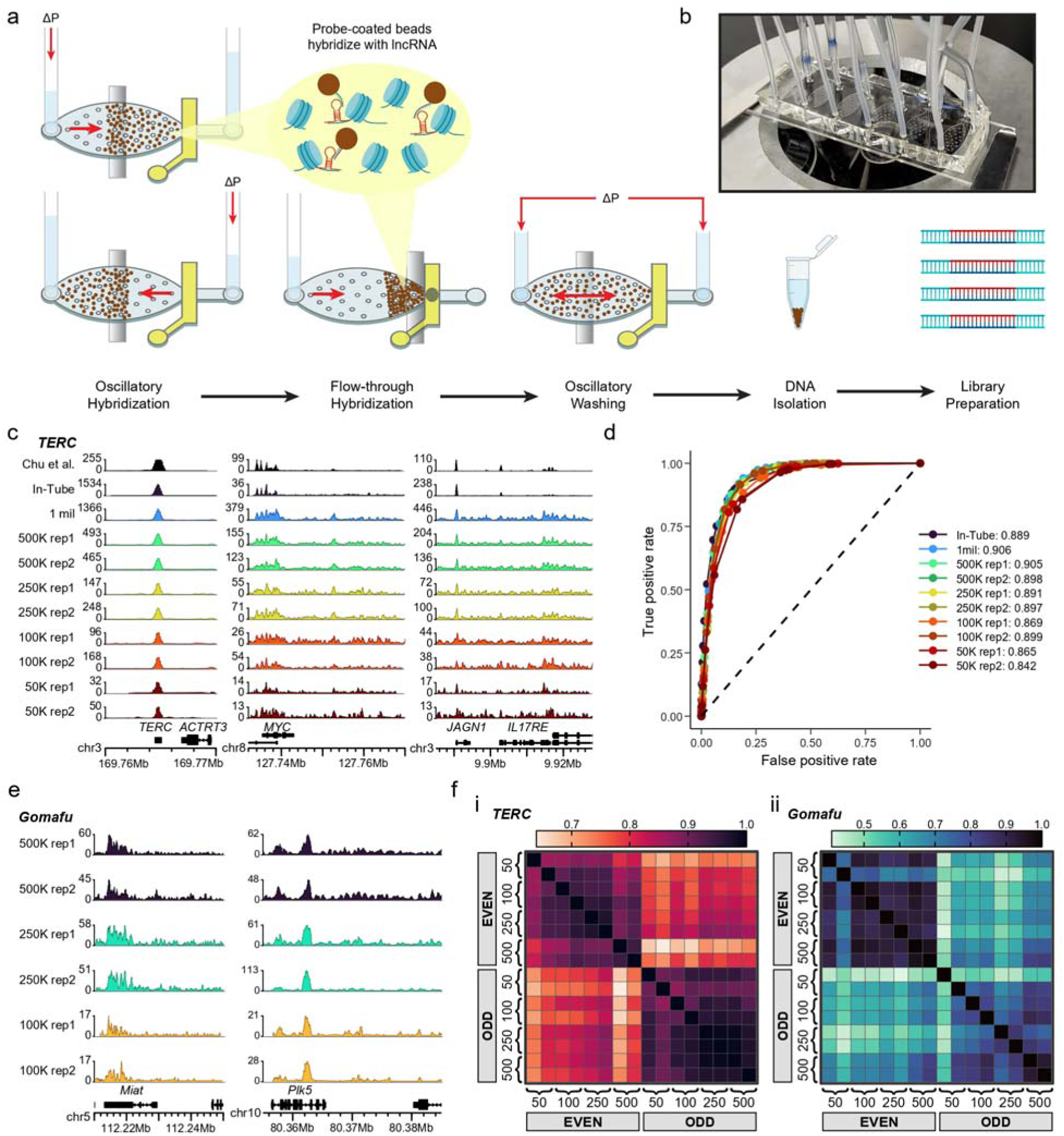
muChIRP-seq workflow, validation against conventional in-tube ChIRP-seq, and reproducibility. (a) muChIRP-seq workflow. A magnet and a sieve valve (controlled by a solenoid valve) facilitate microfluidic device operation. Shaded region above sieve valve indicates control layer is filled with water, depressing the outlet channel and preventing passage of larger particles (magnetic beads). ΔP denotes pressure pulses during oscillatory steps. (b) Glass slide with 6 muChIRP devices. Devices have hybridization tubes in inlet and outlet and a third tube to pressurize the control layer channel. (c) Genome browser tracks showing TERC binding profiles *in HeLa cells* from published in-tube ChIRP-seq data^7^, in-tube ChIRP-seq (20 million cells) generated by us, and muChIRP-seq at decreasing inputs (1 million to 50,000 cells). Tracks show overlapping true peaks, presenting the minimum signal from either even or odd data set. (d) Receiver operating characteristic (ROC) curves comparing muChIRP-seq and in-tube ChIRP-seq to published ChIRP-seq data^7^. Curves reflect performance across different input cell quantities. Area under the curve (AUC) values are shown, preceded by the number of cells used. (e) Gomafu binding in NE-4C cells by muChIRP-seq at decreasing inputs. (f) Pearson correlation of 100-bp binned genomic signal between replicate datasets for inputs ranging from 50,000 to 500,000 cells, for (i) *TERC* binding in HeLa cells and (ii) *Gomafu* binding in NE-4C cells.

### Validation against conventional “in-tube” ChIRP-seq

To benchmark the method, we applied muChIRP-seq to a lncRNA *TERC* (∼450 bp in length)^40^ in HeLa cells and compared our results to published ChIRP-seq data collected using 20 million cells^7^. To control for probe design, we used the same biotinylated tiling oligonucleotide design as in the published study^7^. We optimized specificity and yield of captured ChIRP DNA using muChIRP followed by qPCR to measure fold enrichment at multiple known positive *TERC*-binding loci relative to a negative control (**Supplementary Figure 3**). Optimal fold enrichment and DNA capture were obtained using 100 μL of magnetic beads coated with 20 μL of 10 μM even or odd probes, a 1:1 volume ratio of hybridization buffer to chromatin solution, and five on-device oscillatory washes (see methods). “Even” and “odd” ChIRP-seq datasets were merged and only overlapping signal was considered “true” peaks that identified the genome-wide binding sites^7,8^.

MuChIRP-seq datasets using 1 million to 50,000 input cells were compared against published *TERC* data ^7,8^ and conventional in-tube ChIRP-seq data using 20 million input cells generated by us (**Figure 1c**). Notably, we produced similar binding profiles to the 20-million-cell datasets at key loci using less than 1% of the input material. Genome browser tracks for muChIRP-seq were highly comparable to conventional ChIRP-seq datasets for inputs from 1 million to 250,000 cells. As expected, datasets for 100,000 and 50,000 cells showed lower signal-to-noise ratio and reduced reproducibility compared to datasets with higher input quantities. Defining the published *TERC* data as the gold standard, we quantified agreement of our datasets using a receiver operating characteristics (ROC) curve. MuChIRP-seq datasets from 1 million to 250,000 cells, and one of our two 100,000 cell replicates, reproduced published *TERC* peaks with AUC values (0.899 - 0.906) comparable to that of our in-tube dataset (0.889) (**Figure 1d** and **Supplementary Figure 4).**

### MuChIRP-seq reproducibility

We expanded our studies to *Gomafu* (*Miat*), a lncRNA implicated in schizophrenia^31–33^ with a much longer sequence (∼9-kb in *Mus musculus*^41^ and ∼10-kb in *Homo sapiens*^28–30^) than *TERC*. There were no ChIRP-seq data on *Gomafu* in the literature before this study. We collected replicates at 500,000 to 50,000 cells for *Gomafu* in NE-4C and additional replicates at 100,000 cells for *GOMAFU* in HeLa. Replicate tracks for both *Gomafu* and *TERC* demonstrated strong peaks at known binding loci with high signal-to-noise ratio down to 250,000 cells, with reduced signal-to-noise ratio and reproducibility for 100,000 and 50,000 cells (**Figure 1c** and **1e**, **Supplementary Figures 5-9**). *GOMAFU* and *TERC* were easily distinguishable by their peak profiles, particularly at their transcription loci (**Figure 1c** and **1e, Supplementary Figures 5 and 6**).

We calculated Pearson correlation coefficients among datasets at 100-bp windows across the genome (**Figure 1f**). Correlations among *TERC* datasets in HeLa, ranged from 0.99 to 0.86 between replicates with an overall average cross-dataset correlation of 0.85 ± 0.09, whereas *Gomafu* datasets in NE-4C, ranged from correlation of 0.95 to 0.5 between replicates with an overall average cross-dataset correlation of 0.69 ± 0.15, revealing a steeper decline in data quality with low cell counts in *Gomafu*. We observed that the datasets created using the same probe pool (even or odd) are substantially more correlated than the ones with different probe pools (**Figure 1f**). The average correlations between datasets using same probe pool were 0.92 ± 0.04 for *TERC* and 0.80 ± 0.13 for *Gomafu*, while they were 0.78 ± 0.05 for *TERC* and 0.60 ± 0.10 for *Gomafu* between datasets using different probe pools. Since even and odd probe pools demonstrate similar binding profiles at genomic regions of interest, this suggests that inconsistencies between probe pools occur in off-target regions, underscoring the importance of using separate probe pools and defining high-confidence peaks as those supported by both.

### Cell-type-specific muChIRP-seq using mouse brain tissue samples

The low-input feature of muChIRP-seq makes it uniquely suited for profiling cell-type-specific lncRNA binding using tissue samples. We applied muChIRP-seq to anti-NeuN-stained, FACS-sorted nuclei from mouse cortex to generate cell-type-specific ChIRP-seq data. Two muChIRP-seq replicates were collected on *Terc* binding using each of NeuN+ and NeuN- nuclei samples. Even and odd genome browser tracks showed strong, reproducible peaks at the *Terc* transcription site and consistent signal in *Terc*-associated loci *Myc* and *Tyrobp*^42^ (**Figure 2a**). Genomic distribution analysis showed enrichment of *Terc* true peaks in promoter regions (**Figure 2b**).

**Figure 2.**
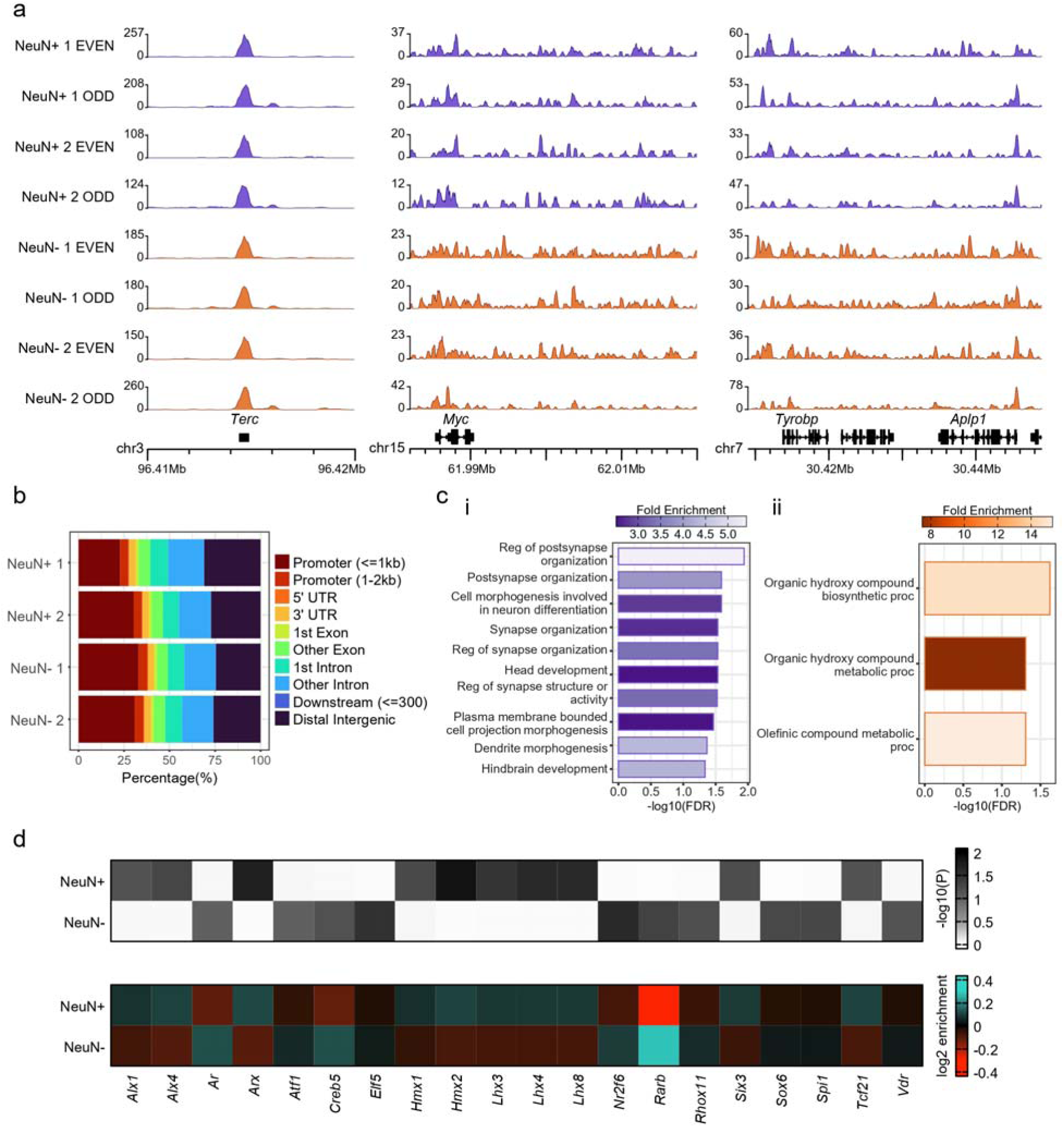
Cell-type-specific muChIRP-seq in mouse cortex. (a) Even and odd genome browser tracks showing *Terc* binding in NeuN+ (neuronal) and NeuN- (glial) nuclei derived from mouse cortex, across replicates. (b) Genomic distribution of true *Terc* binding sites in NeuN+ and NeuN- nuclei. (c) Significantly enriched GO biological processes within stringent cell-type-specific consensus true *Terc* peak sets for (i) NeuN+ and (ii) NeuN- nuclei. (d) TF binary motif analysis of true consensus peaks between replicates for NeuN+ and NeuN- nuclei.

We observed both cell-type-specific binding sites and shared regions between NeuN+ and NeuN- nuclei samples (**Supplementary Figure 10**). We identified a more stringent set of 39 NeuN- specific consensus peaks and 219 NeuN+ specific consensus peaks. These specific consensus peaks appeared in both replicates of each cell type but had no overlaps between cell types in true peaks or individual even/odd datasets. We determined the top ten significantly enriched GO biological processes for each cell-type-specific consensus peak set (**Figure 2c**). GO biological processes exclusively enriched in neuronal (NeuN+) data were primarily related to synapse organization and brain development, suggesting that neuron-specific peaks reflect biologically meaningful differences in *Terc* activity in neurons. In contrast, glial (NeuN-) data presented enriched metabolic process-related terms. These results suggest that while *Terc* binding serves synaptic functions in neurons, its roles in glia are more general.

Lastly, we performed binary transcription factor (TF) motif analysis by comparing the full consensus peak sets for each cell type. These sets comprised all true peaks which appeared in both replicates and were used to identify motifs enriched in one cell type relative to the other (**Figure 2d**). For our analysis, we focused on the top ten motifs by Padj for each cell type. Motifs enriched in neurons included homeo domain factors such as *Hmx1*, *Hmx2*, *Lhx3*, *Lhx4*, *Lhx8*, and *Six3*, which regulate neuronal development and differentiation, consistent with neuronal GO biological processes^43–49^. In contrast, motifs enriched in glia, including *Nr2f6*, *Sox6*, *Spi1*, and *Vdr*, are largely associated with disease states such as cancer and neurodegenerative diseases like Alzheimers^50–54^.

### MuChIRP-seq reveals cell-type-specific lncRNA binding in postmortem human brain

We applied muChIRP-seq to postmortem human prefrontal cortex (PFC) to profile *TERC* and *GOMAFU* binding in disease-relevant neuronal populations. NeuN+ nuclei were profiled from a cohort of schizophrenia subjects and controls. After quality control, we analyzed *GOMAFU* data from three control and four schizophrenia samples, and *TERC* data from four control and four schizophrenia samples. Technical replicates were conducted for control sample CNTRL-24 to confirm muChIRP-seq reproducibility in postmortem PFC (**Supplementary Figure 11**).

True *TERC* peaks were highly concentrated in promoter regions, suggesting that *TERC* may regulate gene expression primarily through interactions at transcription start sites (TSS) (**Figure 3a**). In contrast, *GOMAFU* showed lower enrichment in promoter regions, suggesting a possible regulatory role in intergenic or intronic regions. We performed principal component analysis (PCA) for all true peaks in *GOMAFU* and *TERC* datasets (**Figure 3b**). *TERC* data showed no clear separation between control and schizophrenia, whereas *GOMAFU* data clustered largely by diagnosis. Pearson correlation and hierarchical clustering corroborated these results (**Supplementary Figure 12**). This is consistent with current knowledge of *TERC* function as a telomerase associated lncRNA, predominantly involved in processes such as aging and cancer^55^ and *GOMAFU* playing regulatory roles in schizophrenia^32–34^

**Figure 3.**
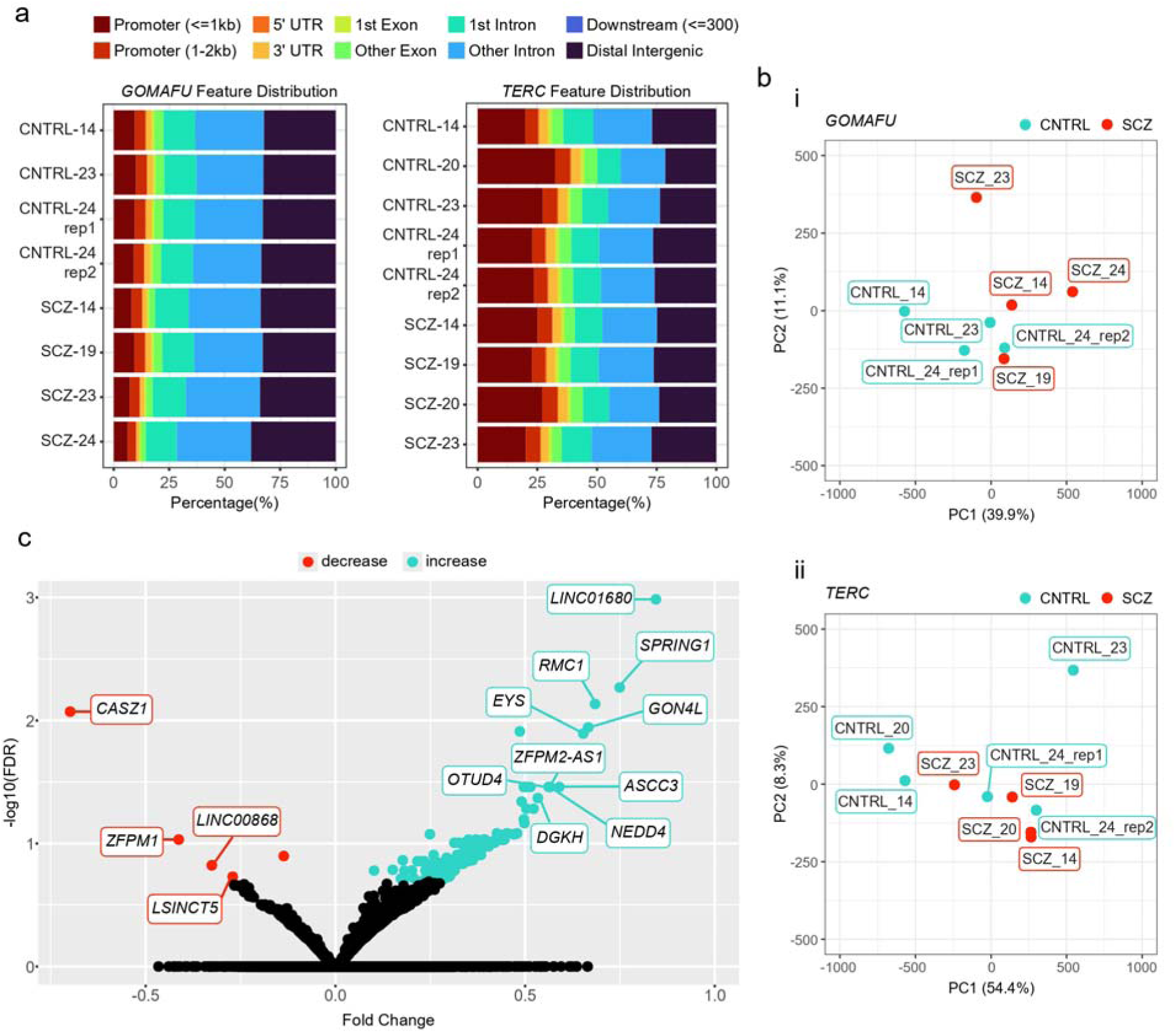
muChIRP-seq profiling of *TERC* and *GOMAFU* in NeuN+ nuclei from postmortem human prefrontal cortex. (a) Genomic distribution of true peak locations for *TERC* and *GOMAFU*. (b) Principal component analysis (PCA) of sample signal across true peak locations for (i) *GOMAFU* and (ii) *TERC*. (c) Volcano plot of significantly differential *GOMAFU* peaks in schizophrenia vs. control. Peaks are separated by increased or decreased fold change in schizophrenia. Selected differential peaks are annotated with gene names.

Differential binding affinity analysis identified 160 *GOMAFU* differential peaks and only one *TERC* differential peak at FDR ≤ 0.2 (**Figure 3c** and **Supplementary Figure 13**). Of the 160 differential *GOMAFU* peaks, 155 showed increased fold change in schizophrenia and only five were decreased, suggesting that elevated *GOMAFU* regulatory action may be characteristic of the disease state. We linked differential *GOMAFU* peaks to their closest genes (158). Among them, *GON4L* showed significantly increased *GOMAFU* binding (**Figure 3c**). *GON4L* has been implicated in autism spectrum disorder^56,57^, and its encoded protein, like *GOMAFU*, localizes to the nucleus^28–35,41,58^. Although a direct link between *GON4L* and *GOMAFU* has not been reported, their shared nuclear localization, involvement in transcriptional regulation, and associations with psychiatric disorder, suggest potential functional interaction. Additional genes showing significant *GOMAFU* binding changes between schizophrenia and control samples included *DGKH*^59^, *NEDD4*^60^, and *CASZ1*^61^, all previously implicated in neurodevelopment or psychiatric disease.

### Profiling data reveal distinct roles for *GOMAFU* and *TERC* in schizophrenia and multimodal analysis suggests coordination among lncRNA binding and other epigenomic modalities

To better understand how lncRNA biology coordinates with other epigenetic modalities and the transcriptomic landscape, we integrated muChIRP-seq data with ChIP-seq datasets on H3K4me3 and H3K27ac as well as RNA-seq data on the same postmortem human brain samples ^36^. For our comparative analysis, we focused on a subset of four schizophrenia subjects and three controls which passed quality control for *GOMAFU*. Within this sample set, we identified 48 differential *TERC* peaks, 160 *GOMAFU* differential peaks, 2,662 differentially expressed genes (RNA-seq),1,402 differential enhancers (H3K27ac), and 10,426 differential promoters (H3K4me3). (**Figure 4a**). Following gene annotation, we examined overlap in associated genes across differential muChIRP-seq, ChIP-seq, and RNA-seq datasets (**Figure 4b**). Among 158 differential *GOMAFU*-associated genes discovered by ChIRP-seq data; 21 were linked to changes in gene expression, 63 to differential promoters, and 11 to differential enhancers. *TERC* and *GOMAFU* target genes showed no overlap. Among the 48 differential *TERC*-associated genes, 8 were linked to gene expression changes, 15 to differential promoters, and 2 to differential enhancers.

**Figure 4.**
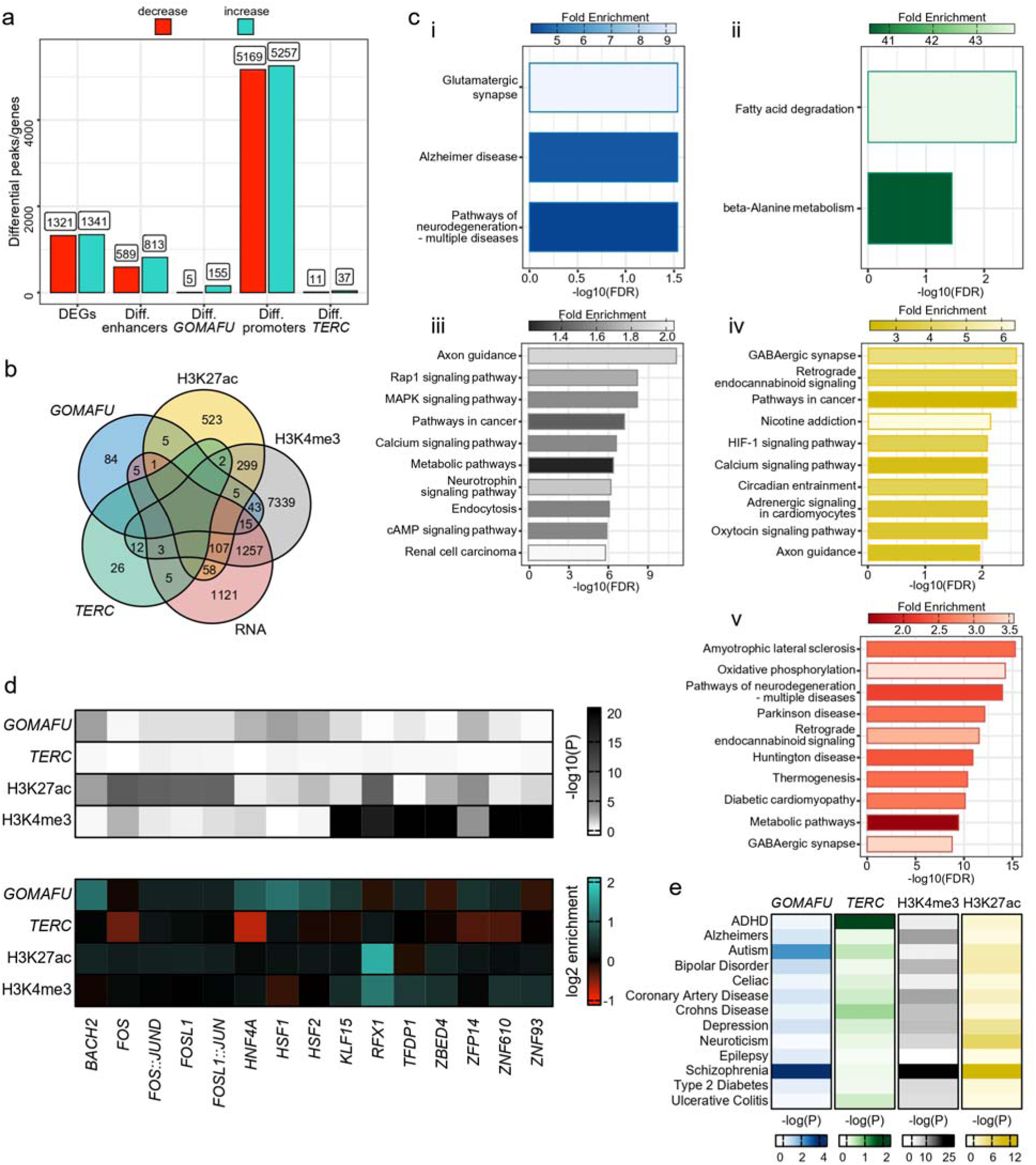
Integrated multiomic analysis reveals different levels of involvement for *GOMAFU* and *TERC* in schizophrenia. NeuN+ nuclei from postmortem human prefrontal cortex samples were used in muChIRP-seq profiling. (a) Significantly differential peaks and genes with increased or decreased enrichment in schizophrenia. Differentially expressed genes (DEGs) identified from RNA-seq data; differential enhancers from H3K27ac data; and differential promoters from H3K4me3 data, restricted to promoter regions defined as ±2 kb from the transcription start site (b) Overlap of differential genes across datasets. Peaks annotated with nearest gene for *GOMAFU*, *TERC*, and H3K4me3, and with published Hi-C data^86^ for H3K27ac. (c) Significantly enriched KEGG pathways within differential genes/peaks for (i) *GOMAFU*, (ii) *TERC*, (ii) H3K4me3, (iv) H3K27ac, and (v) RNA-seq datasets. (d) Transcription factor (TF) motif analysis of differential peaks in *GOMAFU*, *TERC*, H3K27ac, and H3K4me3. The top 5 TFs by adjusted P-value (P_adj_) for *GOMAFU*, H3K4me3, and H3K27ac were included. No motifs had Padj < 1 for *TERC* data. (e) LD score analysis of GWAS disease-associated loci enriched in various differential peak sets.

We identified significantly enriched KEGG Pathways and GO Biological Processes for each differential peak or gene set and compared resulting terms across epigenetic and transcriptomic modalities (**Figure 4c** and **Supplementary Figure 14**). Differentially-bound *GOMAFU* genes were highly enriched for “Glutamatergic synapse” a pathway, which has been implicated in schizophrenia^62^, along with other terms related to neurodegenerative disease. In contrast, enriched pathways in *TERC* showed no clear connection to schizophrenia or other neurodegenerative disorders. Differential promoters marked by H3K4me3 were enriched in multiple signaling-related pathways, such as “*Rap1*”, “*MAPK*”, “Calcium”, “Neurotrophin”, and “cAMP” signaling pathways, all previously implicated in schizophrenia^63–67^. Differential enhancers marked by H3K27ac were also enriched for schizophrenia-related pathways including “GABAergic synapse”, “*HIF-1* signaling”, and “Oxytocin signaling”^68,69^ as well as “Calcium signaling” and “Axon guidance”, which were also enriched in differential promoter data. Differentially expressed genes revealed shared enrichment terms with other datasets including “Pathways of neurodegeneration - multiple diseases” (shared with *GOMAFU*) and “GABAergic synapse” (shared with H3K27ac differential enhancers).

TF motif analysis of differential ChIRP and histone peaks revealed transcription factors potentially involved in schizophrenia-associated epigenomic regulation (**Figure 4d**). Notably, schizophrenia-associated transcription factors such as heat shock factors^70,71^ (*HSF1* and *HSF2*) and members of the *AP-1* complex^72,73^ were among the top motif matches across *GOMAFU* and histone differential peak sets. *HSF1* and *HSF2* were enriched for both *GOMAFU* and H3K27ac. *AP-1* complex members such as *FOS* and *FOSL1* were strongly enriched in H3K27ac; *FOS* was also significant in H3K4me3, whereas *FOS::JUND*, *FOSL1*, and *FOSL1::JUN* motifs were enriched in *GOMAFU* data, suggesting coordinated epigenomic regulation. *TERC* displayed little to no enrichment for motifs significant in other datasets and exhibited negative enrichment for several, including *FOS* and *HNF4A*.

Next, we applied LD score partitioned heritability analysis^74–76^ to test for enrichment of 13 GWAS disease traits across various differential peak sets (**Figure 4e**). Schizophrenia GWAS loci were the most enriched trait in differential H3K4me3, H3K27ac, and *GOMAFU* peak sets, but not in the *TERC* one. In contrast, differential *TERC* showed highest enrichment for ADHD and Crohn’s disease. These results further confirm the lack of schizophrenia-relevant regulatory activity in *TERC*, whereas the strong enrichment of schizophrenia GWAS loci in *GOMAFU* and histone modification differential peaks underscores *GOMAFU*’s potential role in disease-associated chromatin regulation and its coordination with key histone marks.

## Discussion

The lack of robust assays for profiling lncRNA-chromatin interactions in tissue samples has been hindering genome-wide data acquisition that is a prerequisite for improved understanding of lncRNA biology. The development of muChIRP-seq addresses this critical gap by increasing ChIRP DNA collection efficiency in a microfluidic device. The oscillatory hybridization and washing in a microscale chamber promote binding of lncRNA to probes and removal of nonspecific binding, respectively.

We validated muChIRP-seq across multiple lncRNAs (*TERC*, 450 bp; *Gomafu*, 9-10 kb), diverse cell types, and FACS-sorted nuclei from brain tissues, demonstrating its ability to resolve cell-type-specific binding. Integrative analysis of muChIRP data with ChIP-seq data on NeuN+ nuclei isolated from postmortem prefrontal cortex corroborated previous observations on coordination between lncRNA and histone modifications^9,14^ in the context of schizophrenia, and further revealed previously unrecognized lncRNA–chromatin interactions, providing new biological insights into the epigenetic underpinnings of the disease. Our lncRNA profiling data strongly implicate the involvement of *GOMAFU* in schizophrenia pathogenesis while suggesting no involvement of *TERC* in the disease.

MuChIRP-seq is well suited for screening involvement of lncRNAs in diseases at the genome scale, particularly using scarce tissue samples and cell-type-specific subsets. As we demonstrate here, the technology is scalable for screening a myriad of candidate lncRNAs in relevant tissue samples for their potential involvement and roles in a specific disease. Such knowledge is seriously lacking for the vast majority of known lncRNAs. Furthermore, muChIRP-seq data can be readily integrated with other epigenomic and transcriptomic assays for understanding regulatory roles of lncRNAs and their coordination with other epigenetic modalities. Taken together, muChIRP-seq represents a powerful and versatile platform for advancing lncRNA research across various fronts including biomarker discovery, molecular medicine, large-scale functional screening, and basic mechanistic studies of diseases.

## Methods

### Cell culture

HeLa cells (CCL-2; ATCC) were cultured in DMEM (11965092; Gibco) supplemented with 10% fetal bovine serum (FBS; 26140079; Gibco) and 1% penicillin–streptomycin (15140-122; Gibco). NE-4C cells (CRL-2925; ATCC) were cultured in EMEM (30-2003; ATCC) supplemented with 2 mM L-glutamine (G7513; Sigma-Aldrich), 10% FBS, and 1% penicillin–streptomycin. Both cell lines were propagated at 37 °C in a humidified incubator (Symphony, VWR) with 5% CO_2_ and were sub-cultured at ∼80% confluency.

### Mouse brain samples

The animal study was approved by the Institutional Animal Care and Use Committee at Virginia Tech. C57BL/6J mice were purchased from Jackson Laboratory and housed in standard breeding cages at constant temperature of 22 ± 1°C and a 50% relative humidity with 12 h light-dark cycles and with access to food, water ad libitum. 80-week-old male mice were sacrificed by cervical dislocation. Mouse brains were rapidly dissected, frozen on dry ice, and stored at −80°C

### Postmortem human brain samples

Postmortem human brain samples were obtained via autopsies performed at Basque Institute of Legal Medicine, Bilbao, Spain. The study was conducted in accordance with institutional policies and ethical guidelines for postmortem brain studies^77^. The Institutional Review Board (IRB) determined that approval was not needed for this study. Samples were obtained via the opting-out policy, no compensation was provided for tissue donation, and samples were documented with reversible anonymization. Retrospective review of hospital and psychiatric records was used to confirm clinical diagnoses and treatment history. We chose a subset of postmortem brain samples that were profiled for histone modifications and transcriptomes in our previous work^36^, based on availability. They included 5 subjects who met DSM-IV criteria for schizophrenia and 4 controls, all of Caucasian ancestry (**Supplementary Data 1**). Toxicological screening was performed on blood samples taken at the time of death, and on postmortem brain, when possible, to test for the presence of antipsychotics, ethanol, and other drugs. Toxicological studies were performed at the National Institute of Toxicology, Madrid, Spain, using methods including radioimmunoassay, enzymatic immunoassay, high-performance liquid chromatography (HPLC), and gas chromatography-mass spectrometry (GC-MS). The controls were chosen based on negative medical history for neuropsychiatric disorders and drug abuse and negative results in the toxicological screening for all drugs aside from ethanol. Frontal Cortex samples were dissected at autopsy (0.5–1 g tissue) on an ice-cold surface and immediately stored at –80 °C. Schizophrenia samples were classified as antipsychotic-treated or antipsychotic-free based on toxicological screening of blood at the time of death.

### Cell harvest and crosslinking

Confluent cells were recovered by treatment with 0.25% trypsin-EDTA (25200072; Gibco) for 5 min at room temperature. Trypsin was quenched with an equal volume of culture medium, and cells were dislodged by flushing the flask surface and collected. The suspension was centrifuged at 800 g for 4 min, and the medium was aspirated. Roughly 40 million cells were washed once with 30 mL of PBS (pH 7.4; 10010031; Gibco), centrifuged at 800 g for 4 min, and the PBS was aspirated. Cells were crosslinked on an end-to-end rotator for 10 min at room temperature in 40 mL freshly prepared 1% glutaraldehyde (G5882, Sigma-Aldrich). The crosslinking reaction was quenched by adding 10% (v/v) of 1.25 M glycine (G7126; Sigma-Aldrich) into the fixation solution with incubation on an end-to-end rotator for 5 min at room temperature. Crosslinked cells were pelleted by centrifugation at 2,000 g for 5 min, and the supernatant was removed. The pellet was washed with chilled PBS and centrifuged at 2,000 g for 5 min. PBS was removed, and cells were washed again with 2 mL chilled PBS. The suspension was centrifuged at 2,000 g for 3 min at 4 °C. PBS was removed, and cell pellets were stored at –80 °C.

### Nuclei isolation and crosslinking

For homogenization of brain tissue, a piece of tissue (approximately 0.5 cm in diameter) was placed in 3 mL ice-cold nuclei extraction buffer (0.32 M sucrose, 5 mM CaCl2, 3 mM Mg(Ac)2, 0.1 mM EDTA, 10 mM Tris-HCl, and 0.1% Triton X-100), freshly supplemented with 30 μL protease inhibitor cocktail (PIC; P8340; Sigma-Aldrich), 3 μL 100 mM PMSF (P7626; Sigma-Aldrich), 3 μL 1 M DTT(43816; Sigma-Aldrich), and 4.5 μL recombinant RNase inhibitor (RRI; 2313B; Takara Bio). The tissue was homogenized on ice using a Dounce homogenizer (D9063; Sigma-Aldrich), with 15 strokes using loose pestle A followed by 25 strokes with tight pestle B. The homogenate was then filtered through a 40 μm cell strainer (22-363-547; Fisherbrand) and centrifuged at 1,000 g for 10 min at 4 °C. The supernatant was removed, and the pellet was resuspended in a mixture of 500 μL nuclei extraction buffer and 750 μL of 50% iodixanol solution (50% iodixanol, 25 mM KCl, 5 mM MgCl2, and 20 mM Tris-HCl [pH 7.8]). The sample was centrifuged at 10,000 g for 20 min at 4 °C, and the supernatant was aspirated. The pellet was incubated with 300 μL of 2% normal goat serum (50062Z; Thermo Fisher Scientific) in DPBS (14190144; Gibco), freshly supplemented with 3 μL PIC, 0.3 μL 100 mM PMSF, 0.3 μL 1 M DTT, and 0.3 μL RRI for 10 min on ice. Nuclei were resuspended in the same solution of the same volume (∼300 μL) and incubated with 10 μL of 2 ng/μL anti-NeuN antibody conjugated to Alexa Fluor 488 (MAB377X; Sigma-Aldrich) in DPBS on an end-to-end rotator at 4 °C for 1 h, followed by FACS sorting to select (NeuN+) nuclei. The process generates ∼0.3 million neurons from a mouse or postmortem human brain tissue. Multiple pieces of tissue samples were processed if a large quantity of neurons were required. Following FACS, 20 mL PBS, 5 mL 1.8M Sucrose (S0389; Sigma-Aldrich), 250 μL 1M CaCl_2_ (21097; Sigma-Aldrich), and 75 μL 1M Mg(Ac)_2_ (M5661; Sigma-Aldrich) were added into 5 mL of nuclei suspension (containing ∼1 million sorted nuclei). The sample was incubated on ice for 15 min and then centrifuged at 1,800 g for 15 min at 4 °C, and the supernatant was discarded. The pellet was resuspended in 960 μL PBS. Then, 40 μL of 25% glutaraldehyde was added, resulting in a 1% glutaraldehyde crosslinking reaction. The sample was crosslinked at room temperature on an end-to-end rotator for 10 min. The reaction was quenched by adding 10% (v/v) of 1.25 M glycine for 5 min at room temperature on an end-to-end rotator. Crosslinked nuclei were pelleted by centrifugation at 2,000 g for 5 min, washed with chilled PBS, and centrifuged again. Nuclei were washed once more with 50 μL chilled PBS and centrifuged at 2,000 g for 3 min at 4 °C. Nuclei pellets were stored at −80 °C. The crosslinking process can be scaled up when a larger quantity of nuclei are required.

### Lysis and sonication

1 million crosslinked cells or nuclei were resuspended in 50 μL lysis buffer (50 mM Tris-Cl [pH 7.0], 10 mM EDTA, and 1% SDS), supplemented with 1% (v/v) each of PIC, 100 mM PMSF, and RRI. The suspension was sonicated using a Bioruptor Plus (UCD-300, Diagenode) with a 4 °C water bath, set to HIGH (30 s on, 45 s off pulse interval) for 30-min cycles. Following two sonication cycles (1 h), 2 μL of lysate was transferred to a 1.5 mL Eppendorf tube and 93 μL of Proteinase K buffer (100mM NaCl, 10 mM Tris-Cl [pH 8.0], 1 mM EDTA, and 0.5% SDS) and 5 μL Proteinase K (PK; P2308; Sigma-Aldrich) were added and sample was briefly vortexed and placed on a Thermal Shaker for 45 min at 50 °C and 1,000 rpm. Sample was briefly centrifuged and 100 μL phenol chloroform (P3803; Sigma-Aldrich) was added. Solution was vortexed vigorously for 10 s and then centrifuged at 16,100 g for 5 min at room temperature. The top liquid phase was carefully extracted and 480 μL 100% ethanol, 60 μL 5 mM ammonium acetate (A2706; Sigma-Aldrich), and 6 μL 5 mg/mL glycogen (AM9510; Invitrogen) were added. The solution was incubated for 2 h at −80 °C. Following incubation, sample was centrifuged at 16,100 g for 10 min at 4 °C. Supernatant was discarded and the DNA pellet retained. The pellet was washed with 500 μL ice-cold 70% ethanol then centrifuged at 16,100 g for 5 min at 4 °C. Ethanol was discarded, and sample was centrifuged at 16,100 g for 30 s at 4 °C. Remaining ethanol was removed from sample and the pellet was air-dried in a biosafety cabinet for 10 min. Dried pellet was resuspended in 10 μL low EDTA TE. Chromatin size was measured via TapeStation (Agilent) following manufacturer protocol. Sonication was considered complete if the majority of chromatin fragments measured 100–500 bp in length (**Supplementary Figure 2**). Sonication and size check steps were repeated on remaining lysate as necessary to achieve the target fragment size distribution (2-6 cycles total). Following sonication, remaining lysate was centrifuged at 16,000 g for 10 min at 4 °C. The supernatant was retained, and the pellet was discarded. Sonicated chromatin was stored at −80 °C prior to use in a muChIRP experiment.

### Probe design

Biotinylated tiling oligonucleotides for human *TERC* matched those used in published ChIRP-seq data^7^.17 probes for mouse *TERC* were designed from RefSeq transcript NR_001579.1 using the same technique. Two sets of 48 probes for human and mouse *Gomafu* were designed with the LGC Biosearch Technologies ChIRP-seq probe designer, following parameters from published ChIRP-seq protocols^7,8^ (**Supplementary Data 2**). Transcript sequences NR_003491.4 (human) and NR_033657.1 (mouse), both from RefSeq, were used as references.

### Preparation of probe-coated beads

Dried oligonucleotide probes (LGC Biosearch Technologies) were resuspended following manufacturer’s protocol to a concentration of 100 μM for each probe in low-EDTA TE buffer (IDT) yielding 500 μL of each individual probe solution. 10 μL each of the even-numbered probes were combined to create a 100 μM even probe set pool. 10 μL each of the odd-numbered probes to create a 100 μM odd probe set pool. 100 uM probe set pools and remaining 100 μM individual probe solutions were stored long term at −20 °C. 100 μM even and odd probe sets pools were further diluted to 10 μM in low-EDTA TE for short-term storage at 4 °C for up to a month. 100 μL of Dynabeads Streptavidin MyOne C1 magnetic beads (65001; Invitrogen) were prepared for each probe set. Beads were washed twice with 170 μL lysis buffer. They were then resuspended in 280 μL lysis buffer containing 20 μL of 10 μM even or odd probe set pool. Beads were incubated on an end-to-end rotator at 37 °C overnight. Prior to use in a muChIRP experiment, beads were washed twice with 170 μL lysis buffer, resuspended in 100 μL hybridization buffer (750 mM NaCl, 1% SDS, 50 mM Tris-Cl [pH 7.0], 1 mM EDTA, and 15% formamide), and freshly supplemented with 1% (v/v) each of 100 mM PMSF, PIC, and RRI.

### Fabrication of microfluidic devices

The muChIRP device was fabricated using multi-layer soft lithography^38,39^. Two photomasks (one for the control layer and one for the fluidic layer) were designed using LayoutEditor and printed on high-resolution transparencies (10,000 dpi; Fineline Imaging) (**Supplementary Figure 1b** and **1c**). The fluidic layer design consists of inlet and outlet channels connected to an elliptical chamber (6 mm × 12 mm). Pillars were positioned within the chamber to prevent collapse. The control layer consists of a 4 mm × 1 mm channel, which is connected by a narrow channel to an inlet. To fabricate the fluidic master, SU-2075 (MicroChem) was spin-coated on a 3-inch silicon wafer (University Wafers) at 500 rpm for 10 s, followed by 3,250 rpm for 30 s to yield a 72-μm-thick layer. For the control layer master, SU-2075 was spun at 500 rpm for 10 s, then 1,900 rpm for 30 s, yielding a 120-μm control layer. The wafers were soft-baked at 95□°C for 8 min, then covered with the corresponding photomask and exposed to UV light for 17 s. The wafers were then baked at 95 °C for 8 min and developed in SU-8 developer (Kayaku Advanced Materials Inc.) for 10 min. Wafers were then washed with acetone, isopropanol, and DI water, then fully dried and hard baked at 150 °C for 15 min. The two-layer muChIRP devices were fabricated out of poly(dimethylsiloxane) (PDMS) (RTV 615; Momentive) using multi-layer soft lithography techniques^38,39^. The control layer was fabricated using a prepolymer mass ratio of 20:1 for A:B, which was spun onto the wafer at 500 rpm for 10 s, followed by 1,100 rpm for 30 s. The fluidic layer was fabricated using a prepolymer mass ratio of 5:1 for A:B, which was poured directly onto the fluidic master in a petri dish. Both layers were partially cured at 80 °C for 12 min. At this point, the fluidic layer was peeled from its master and the inlet and outlet holes were punched using a 2 mm puncher (Harris Uni-Core). The fluidic layer was then adhered to the top of the control layer so that the alignment marks in their respective designs lined up. The two-layer device was baked at 80 °C for 2 h, after which the device was peeled from the control master and the control layer inlets were punched. The PDMS slabs were adhered to precleaned glass slides using a plasma cleaner (PDC-32G; Harrick Plasma), followed by a final 80 °C bake for 4 h.

### Operational setup for muChIRP

Reagents were introduced to the device through 80 cm of perfluoroalkoxy (PFA) loading tubing (1622L, ID: 0.02 in., OD: 0.0625 in.; IDEX Health & Science) attached to a 1 mL syringe via Luer Lock and fitting, with flow driven by a syringe pump (Fusion 400; Chemyx) (**Supplementary Figure 1a**). Short tubing assemblies consisting of 6 cm of C-flex clear tubing connected to 3 cm of PFA tubing were used to actuate the control valve with high pressure water and to house buffer during oscillatory washing. Long tubing assemblies consisting of 6 cm of C-flex clear tubing connected to 45 cm of PFA tubing were used for sample manipulation during oscillatory hybridization. The control valve was actuated by a solenoid valve (18801003-12V; ASCO Scientific) connected to a compressed air source^39^. During oscillatory hybridization and washing steps, the inlet and outlet of the device were each connected to one solenoid valve in a two-set solenoid manifold. A data acquisition card (NI SCB-68; National Instruments) enabled control valve manipulation and automation of oscillatory hybridization and washing steps via LabVIEW (National Instruments).

### MuChIRP

Before running muChIRP, the control channel was attached to a solenoid valve with a short tubing assembly and filled with pressurized water until any bubbles were eliminated. Next, the control valve was opened (no pressure exerted in the control layer), and the microfluidic device was flushed with hybridization buffer via loading tubing assembly attached to the inlet. Flow was driven at 200 μL/min by syringe pump until air was completely driven from the device. Following this, 100 μL probe-coated beads (even or odd) were pipetted into the device, guided by a magnet placed at the inlet, and the control valve was closed. A magnet was adhered to the bottom of the device so that the beads were held in the center of the chamber during following steps (**Figure 1a**). 2.5 - 50 μL of sonicated chromatin solution (from 50,000 - 1 million cells/nuclei, respectively) was diluted 1:1 by volume with hybridization buffer, and the resulting solution was freshly supplemented with 1% (v/v) PIC, 100 mM PMSF, and RRI. A long tubing assembly was attached to the outlet of the device and to one solenoid valve in a two-set solenoid manifold. The full volume of sonicated chromatin in hybridization buffer solution (5 - 100 μL) was drawn into loading tubing via attached syringe and injected into the device at 20 μL/min, flowing into the long tubing assembly at the outlet. Following full sample injection, the control valve was closed and loading tubing assembly was removed from the inlet. A second long tubing assembly was connected to the second solenoid valve in the two-set manifold and to the device inlet. The valve was opened to allow the chromatin solution to flow through the device, into the long tubing at the inlet. Once the liquid levels in the long tubing attached to the inlet and outlet were equal, oscillatory hybridization was initiated. Pressure was set to 0.5 psi, and pulse duration was set from 1-20 s for 5 - 100 μL of chromatin to ensure that all the volume moved through the device during each pair of pulses. The total duration was 4 h, and the process was run unattended via LabVIEW. After the 4-h oscillatory hybridization, the full chromatin solution was directed into the long outlet tubing using LabVIEW to pressurize the long inlet tubing until no sample remained. Next the control valve was closed, and the inlet tubing was removed. The loading tubing assembly was inserted into the inlet, and the control valve was opened. The sample solution was drawn into the tubing via an attached syringe, until no solution remained in the outlet tubing, taking care to avoid air entry into the device. The magnet was placed flush with the closed control valve so that the magnetic beads formed a packed bed. The solution was injected into the device at 1.5 μL/min, until the full volume reached the inlet. After hybridization, the magnet was placed at the center of the device and oscillatory washing was conducted. Two short tubing assemblies were filled with 50 μL of washing buffer (2× NaCl, 2× SSC, and 0.5% SDS) each, freshly supplemented with 1% (v/v) 100 mM PMSF. Tubing assemblies were attached to the inlet and outlet of the device, and to each solenoid valve in a two-set solenoid manifold. Alternating pressure pulses at 1 psi were applied at 0.3 second intervals to the inlet and outlet tubing assemblies for 7.5 min cycles using LabVIEW. This process was repeated with fresh washing buffer for a total of five cycles. Lastly, the streptavidin magnetic beads were flushed from the device at 200 μL/min using remaining hybridization buffer in loading tubing assembly and collected in a 1.5 mL Eppendorf tube.

### DNA isolation

The tube containing streptavidin magnetic beads with bound chromatin was placed in a magnetic rack and the supernatant was removed. The beads were resuspended in 100 μL elution buffer (50 mM NaHCO_3_ and 1% SDS) supplemented with 0.32 μL RNase H (2150B, Takara Bio) and 0.2 μL RNase A (19101, Qiagen). Suspension was incubated on a Thermal Shaker for 30 min at 37 °C and 1,000 rpm. Beads were returned to the magnetic rack, and the supernatant was added to a fresh 1.5 mL Eppendorf tube. Beads were resuspended in 100 μL supplemented elution buffer and incubated on the Thermal Shaker for an additional 30 min at 37 °C and 1,000 rpm. Supernatant was removed from the magnetic beads and added to the same 1.5 mL Eppendorf tube. 10 μL Proteinase K (PK; P2308; Sigma-Aldrich) was added to the 200 μL solution and the sample was placed on a Thermal Shaker for 45 min at 50 °C and 1,000 rpm. Sample was briefly centrifuged and 200 μL phenol chloroform (P3803; Sigma-Aldrich) was added. Solution was vortexed vigorously for 10 s and then centrifuged at 16,100 g for 5 min at room temperature. The top liquid phase was carefully extracted and 960 μL 100% ethanol, 120 μL 5 mM ammonium acetate (A2706; Sigma-Aldrich), and 6 μL 5 mg/mL glycogen (AM9510; Invitrogen) were added. The solution was incubated overnight at −80 °C. Following overnight incubation, samples were centrifuged at 16,100 g for 10 min at 4 °C. Supernatant was discarded and the DNA pellet retained. The pellet was washed with 1 mL ice-cold 70% ethanol then centrifuged at 16,100 g for 5 min at 4 °C. Ethanol was discarded, and sample was centrifuged at 16,100 g for 30 s at 4 °C. Remaining ethanol was removed from sample, and the pellet was air-dried in a biosafety cabinet for 10 min. Dried pellet was resuspended in 5 μL low EDTA TE and stored at −20 °C.

### Construction of sequencing libraries

Libraries from human PFC, mouse cortex and *GOMAFU* in HeLa cells were prepared using the Takara DNA Thruplex Kit, while remaining cell line libraries were prepared with the IDT ACCEL-NGS 2S DNA Library Kit, following manufacturer protocols. Library amplification was performed by real-time qPCR (1855200; Bio-Rad) following the corresponding PCR protocol for each kit. Amplification was stopped once fluorescence increased by 1,500 RFU above baseline. Samples were purified using SPRI magnetic beads (B23317; Beckman Coulter) and the fragment size was measured using the Agilent TapeStation. The KAPA Library Quantification Kit (Roche Diagnostics) was used according to the manufacturer’s protocol to measure library concentration. Libraries were pooled such that approximately 50 million raw reads were allocated to each sample and sequenced on a NovaSeq X instrument to generate 150-bp paired-end reads.

### MuChIRP-qPCR

Real-time qPCR was performed using iQ SYBR Green Supermix (1708886; Bio-Rad) and custom primers targeting known lncRNA-binding regions (IDT) to confirm enrichment of ChIRP DNA at specific loci. Assays for all lncRNAs and tissue types were performed using the following PCR protocol: 95□°C for 10 min followed by 40 cycles of 95□°C for 15 s, 58°C for 40 s, and 72□°C for 30 s. “Percent of input” was calculated for both positive and negative loci, and relative fold enrichment was derived from these values as follows:

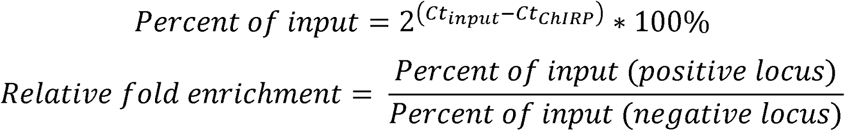

Where Ct_input_ and Ct_ChIRP_ refer to the threshold cycle values of the input and ChIRP DNA, respectively (**Supplementary Figure 3**).

### ChIRP-seq data processing

ChIRP-seq and corresponding input data were aligned to the mouse genome (mm10) or human genome (hg38) using Bowtie 2^78^ in paired-end mode. Peaks were called using MACS2^79^ broad peak calling with a q-value cutoff of 0.05, broad cutoff of 0.1, max-gap= 120 and min-length= 100. True coverage tracks were generated for each even–odd ChIRP-seq pair by taking the minimum coverage between even and odd, at each base pair of the genome as described in Chu et al^7,8^. Genomic coverage tracks were visualized using the karyoploteR package^80^. Unless otherwise specified, true peaks were defined as MACS2 peaks from even and odd datasets that either overlapped by at least 1 bp or were directly adjacent (bookended). Overlapping/bookended peaks were merged into a single peak spanning the combined region (**Supplementary Data 3**). Based on our assessment of cell line ChIRP-seq data and observation of associated quality markers, we set a quality control filter of unique reads > 3,000,000 and true peaks > 8,000 for muChIRP tissue data.

### ChIP-seq data processing

ChIP-seq data were aligned to the human genome (hg38) using Bowtie 2^78^ in single-end mode. Peaks were called using MACS2^79^ with a q-value cutoff of 0.05 for narrow peaks for H3K4me3 data. Only H3K4me3 peaks that overlapped promoter regions (defined as ±2 kb from the transcription start site) by at least 1 bp were retained for downstream analysis. Peaks were called using MACS2^79^ broad peak calling with a q-value cutoff of 0.05, broad cutoff of 0.1, max-gap = 120 and min-length= 100 for H3K27ac data.

### RNA-seq data processing

RNA-seq data was aligned to the human genome (hg38) using HISAT2^81^. FeatureCounts^82^ was used to obtain gene-level read counts.

### Construction of receiver operating characteristic (ROC) curves

ROC curves were used to compare the performance of muChIRP-seq at different starting cell quantities to previously published ChIRP-seq data for *TERC* binding in HeLa cells created by conventional protocol^7^ (**Figure 1d**). We focused on “true peaks” in promoter regions (defined as ±2 kb from the transcription start site). The published ChIRP-seq data for *TERC* in HeLa cells^7^ served as the gold standard. Promoter regions overlapping with a true peak by at least 1 bp were defined as gold standard positives, while promoter regions lacking a true peak were defined as gold standard negatives. When generating curves for our data, true positives (TPs) were promoter region peaks matched to gold standard positives; true negatives (TNs) were empty promoter regions matched to gold standard negatives; false positives (FPs) were peaks not matched to gold standard positives; and false negatives (FN) were empty promoter regions containing a gold standard peak. True positive rate (TPR) was defined as TP/(TP+FN) and false positive rate (FPR) was defined as FP/(FP+TN). ROC curves were generated by varying the MACS2 q-value cutoff. The area under the curve (AUC) was calculated for each ROC curve.

### Correlation analysis

Pearson correlation analysis of cell line data was performed by looking at signal across 100-bp windows of the genome. Correlation of binned signal between datasets was calculated using *cor()* from base R and was plotted using *ggplot2* (**Figure 1f**). For human PFC data, consensus peaks were defined as the full set of true peaks from all samples. Overlapping peaks between even and odd datasets were identified using the *dba.peakset* function in DiffBind^83^. Read counts were obtained using *dba.count*, with the *summits* parameter set to half the average consensus peak width (*summits* = 400 for *TERC*; *summits* = 350 for *GOMAFU*). True peak counts were the minimum counts at each location between even and odd datasets. Correlations were calculated using *dba.plotHeatmap*, and hierarchical clustering and visualization were performed using the *ComplexHeatmap* R package (**Supplementary Figure 12**).

### Principal component analysis (PCA)

Principal component analysis was performed on the full set of true peaks across all samples. Overlapping peaks between even and odd datasets were identified using the *dba.peakset* function in DiffBind^83^. Read counts were obtained using *dba.count*, with the *summits* parameter set to half the average consensus peak width (*summits* = 400 for *TERC*; *summits* = 350 for *GOMAFU*). True peak counts were the minimum counts between even and odd datasets at each peak location. Peak locations with zero variance across samples were removed prior to PCA. PCA was performed using *prcomp()* from the *stats* R package. Sample values for the top two principal components were plotted were plotted using *ggplot2* (**Figure 3b**).

### Differential analysis of postmortem human PFC data

Differential analysis of ChIP-seq and ChIRP-seq data was performed using DiffBind^83^. True peaks for ChIRP-seq were identified using the *dba.peakset* function in DiffBind. Consensus peaks were defined as true peaks which appeared in at least two samples within the same condition (schizophrenia or control). Consensus peaks for ChIP-seq data were defined as peaks that appeared in both replicates and in at least two samples in the same condition. Read counts at each consensus peak were found using the *dba.count* function with the *summits* parameter set to half the average peak width (950 for *TERC*, 550 for *GOMAFU*, 1000 for H3K4me3, and 1500 for H3K27ac). Differential analysis was performed using *dba.analyze* with the *method* set to DESeq2 and its default normalization strategy. Peaks with FDR < 0.2 were considered significantly differential. Differential analysis of RNA-seq data was performed using DESeq2^84^, with gene counts obtained from featureCounts^82^. Genes with fewer than 20 reads in more than 50% of the samples were removed prior to DeSeq2 analysis. Age of patients was regressed out of the model after principal component analysis identified it as being highly correlated with multiple of the top 6 principal components. Default normalization parameters were used, and genes with FDR < 0.2 were considered significantly differential. Differential peaks for ChIRP-seq and H3K4me3 were annotated using ChIPseeker^85^. H3K27ac differential enhancers were annotated using published Hi-C data from neuronal nuclei in human cortex^86^, where possible. Remaining differential H3K27ac peaks were annotated using ChIPseeker (**Supplementary Data 4**).

### GO term analysis & GWAS LD score analysis

Analysis with shinyGO^87^ was performed on significantly differentially expressed genes and genes associated with differential peaks. Significantly enriched GO biological processes and KEGG pathways were defined as those with FDR < 0.05 (**Figure 2c**, **4c**, **Supplementary Figure 14** and **Supplementary Data 5**). GWAS trait enrichment was calculated using LD Score partitioned heritability with default parameters^74–76^ (**Figure 4e**). Summary statistics for each trait were obtained from the 1000 Genomes European population set^88^.

### Transcription factor motif analysis

Transcription factor (TF) motif analysis was performed using monaLisa^89^. Binary motif analysis of mouse cortex data was performed by comparing neuron and glia consensus peaks (defined as all true peaks which appeared in both replicates) using motifs obtained from the JASPAR 2024 *Mus musculus* CORE collection^90^ (**Figure 2d**). Single set motif analysis was performed on each of the sets of differential peaks obtained from postmortem prefrontal cortex samples (**Figure 4d**). TF motifs were obtained from JASPAR 2024, using the *Homo sapiens* CORE collection^90^. TFs of interest were defined as the top ten motifs by P_adj_ for each mouse peak set and the top five for each human prefrontal cortex differential peak set. In the case where two motifs had the same P_adj_, the motif with higher enrichment was chosen.

## Data availability

Published ChIP-seq and RNA-seq data on the postmortem brain samples used in this study were obtained from dbGaP accession number phs002487.v1.p1^36^. Published ChIRP-seq data was obtained from GEO accession number GSE31332^7^. The muChIRP-seq data discussed in this publication have been deposited in NCBI’s Gene Expression Omnibus and are accessible through GEO Series accession number GSE316123 (https://www.ncbi.nlm.nih.gov/geo/query/acc.cgi?&acc=GSE316123).

## Code availability

All codes are available at GitHub (https://github.com/changlulab/Catalano_et_al_2026).

## Acknowledgments

This work was supported by the United States National Institutes of Health grants R01GM141096 (C.L. and Z.B.C.), R01DA056187 (C.L.), R01MH084894 (J.G.-M.), R35HL171550 (Z.B.C.) and by the Basque Government grant IT-1920/26 (J.M.)

## Author contributions

C.L. and Z.B.-C. conceived the project. C.L., J.A.C., Y.-P.H., and Z.L. designed the protocol. J.A.C. conducted all ChIRP-seq experiments and data analysis. G.L. carried out the mouse work. J.M. and J.G.-M. provided postmortem human brain samples and expertise on schizophrenia. J.A.C. and C.L. wrote the manuscript. All authors proofread the manuscript and provided feedback.

