## Supplementary Information for "Microfluidic low-input profiling reveals lncRNA roles in disease"

### Supplementary Figures

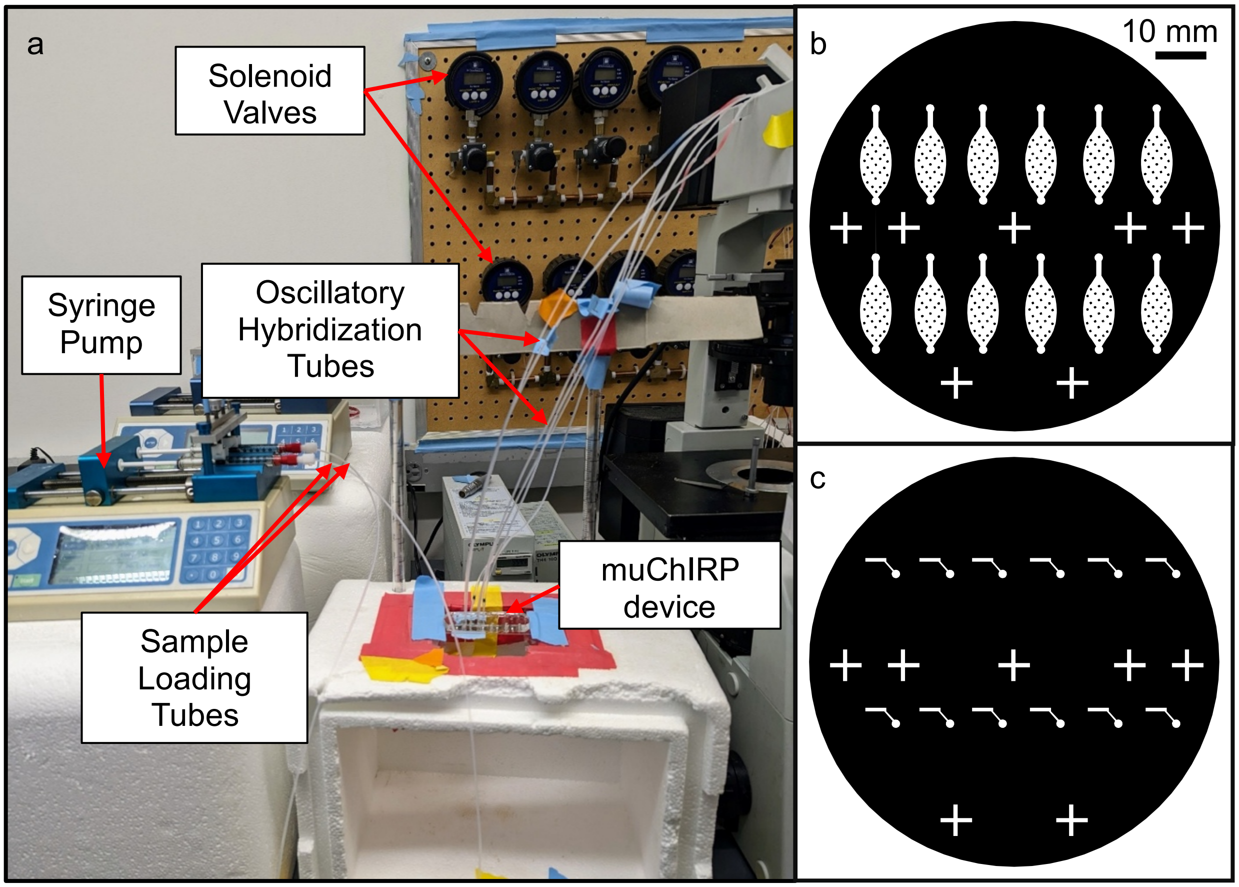


**Supplementary Figure 1. muChIRP microfluidic device set up. (a)** Labeled photograph of the muChIRP on-device experimental setup. **(b)** muChIRP microfluidic device fluidic layer photomask design. The eliptical chamber is approximately 6 mm × 12 mm and connects to an inlet and outlet channel; pillars are positioned to prevent collapse. **(c)** Photomask design for the control layer. A 4 mm × 1 mm channel connects via a narrow diagonal channel to an inlet. Plus marks are used for alignment of the control and fluidic layers. Scale bars (b,c), 10 mm.

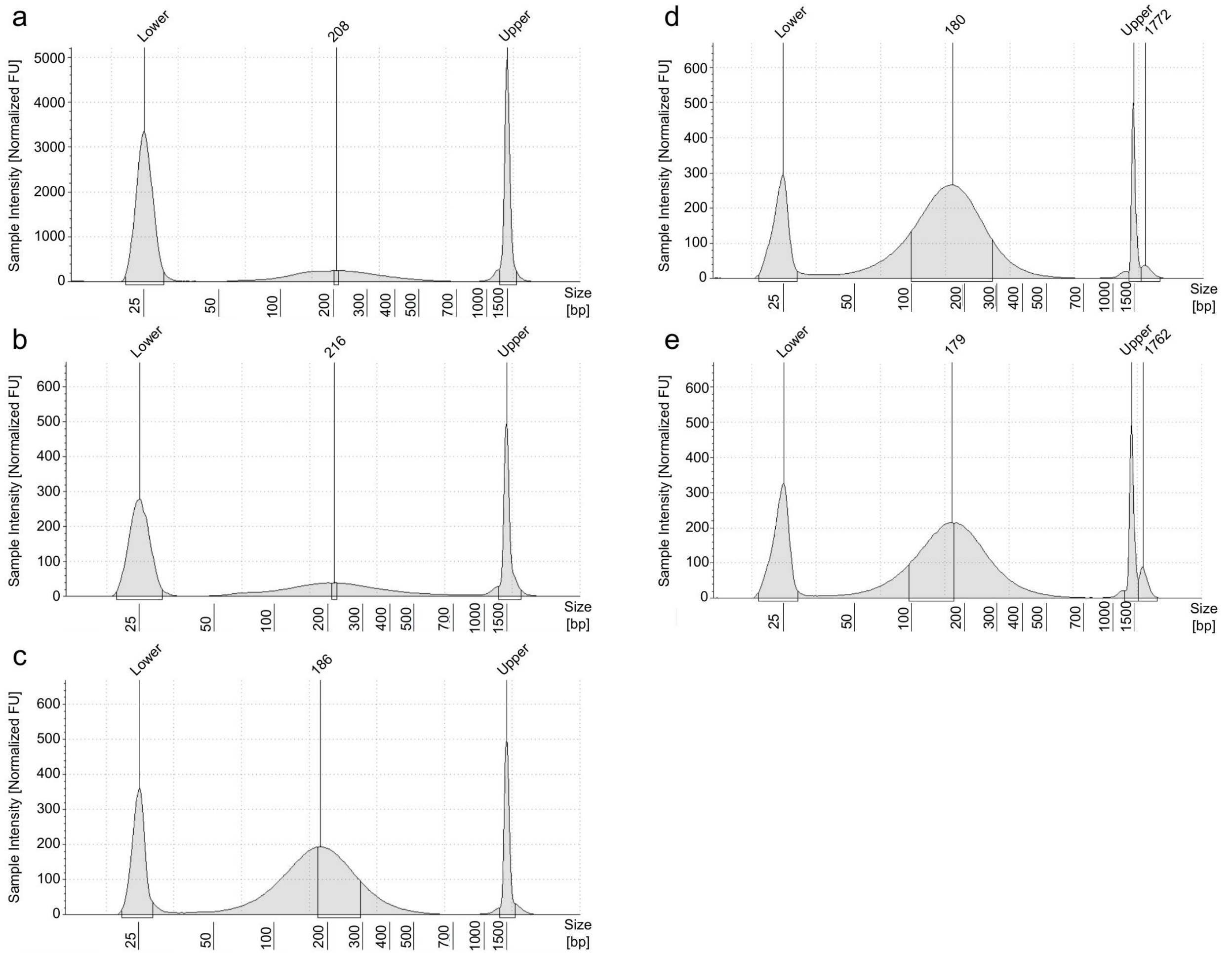


**Supplementary Figure 2. Chromatin fragment size distributions following sonication.** TapeStation electropherograms of sonicated chromatin input from (a) HeLa cells, (b) NE-4C cells, (c) NeuN+ nuclei from postmortem human brain tissue (sample CNTRL-24), (d) NeuN+ nuclei from mouse cortex, and (e) NeuN- nuclei from mouse cortex. Chromatin was sheared to a target size range of 100–500 bp prior to muChIRP.

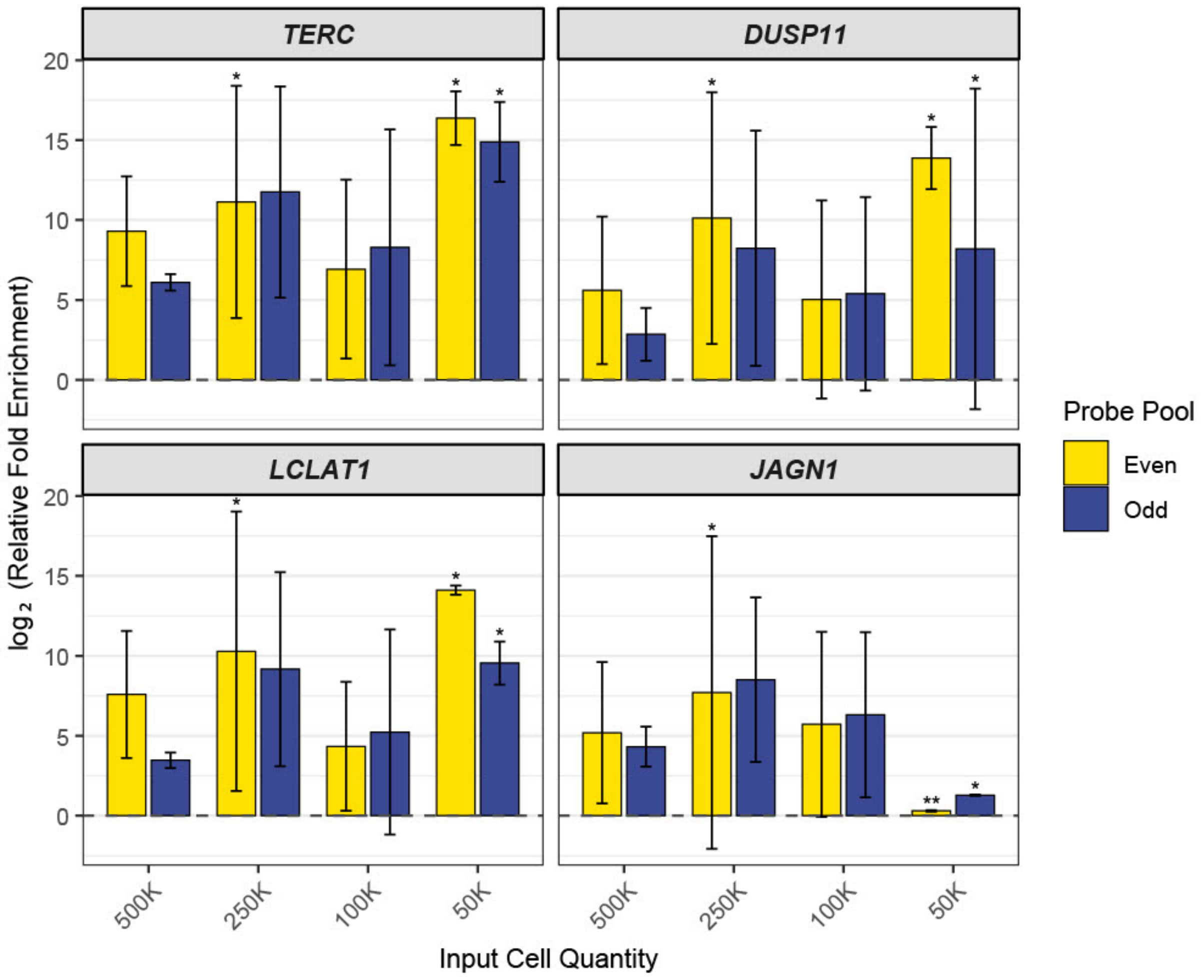


**Supplementary Figure 3. muChIRP-qPCR enrichment at *TERC*-binding loci in HeLa cells.** Relative fold enrichment was calculated as described in Methods, using *AFM* as the negative control locus. Bars represent the mean $\log_2$(relative fold enrichment) from two technical replicates. Error bars represent standard deviation between replicates
* indicates that the negative control locus did not amplify in one or more replicates, and enrichment was calculated by setting Cq=42 for the negative locus.
** indicates that both the negative and positive loci failed to amplify in one or more replicates, and Cq=42 was used for both.

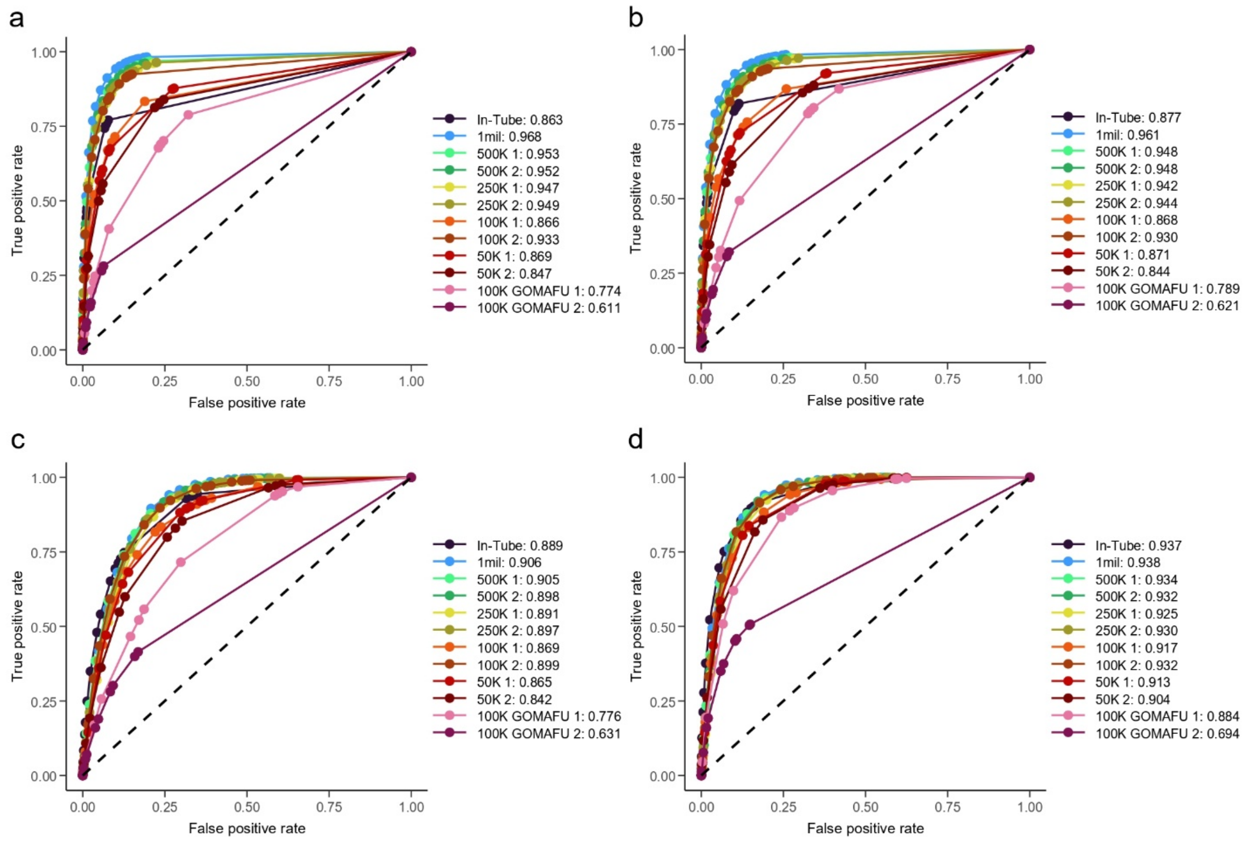


**Supplementary Figure 4. *TERC* in HeLa cells ROC Curves.** Receiver operating characteristic (ROC) curves comparing muChIRP-seq and our in-tube ChIRP-seq to published ChIRP-seq data[7]. Curves reflect performance across different input cell quantities. Area under the curve (AUC) values are shown, preceded by the number of cells used. Each panel compares performance across different types of genomic regions: **(a)** 100-bp bins **(b)** 1000-bp bins **(c)** promoter regions (±2 kb from transcription start sites) **(d)** gene bodies.

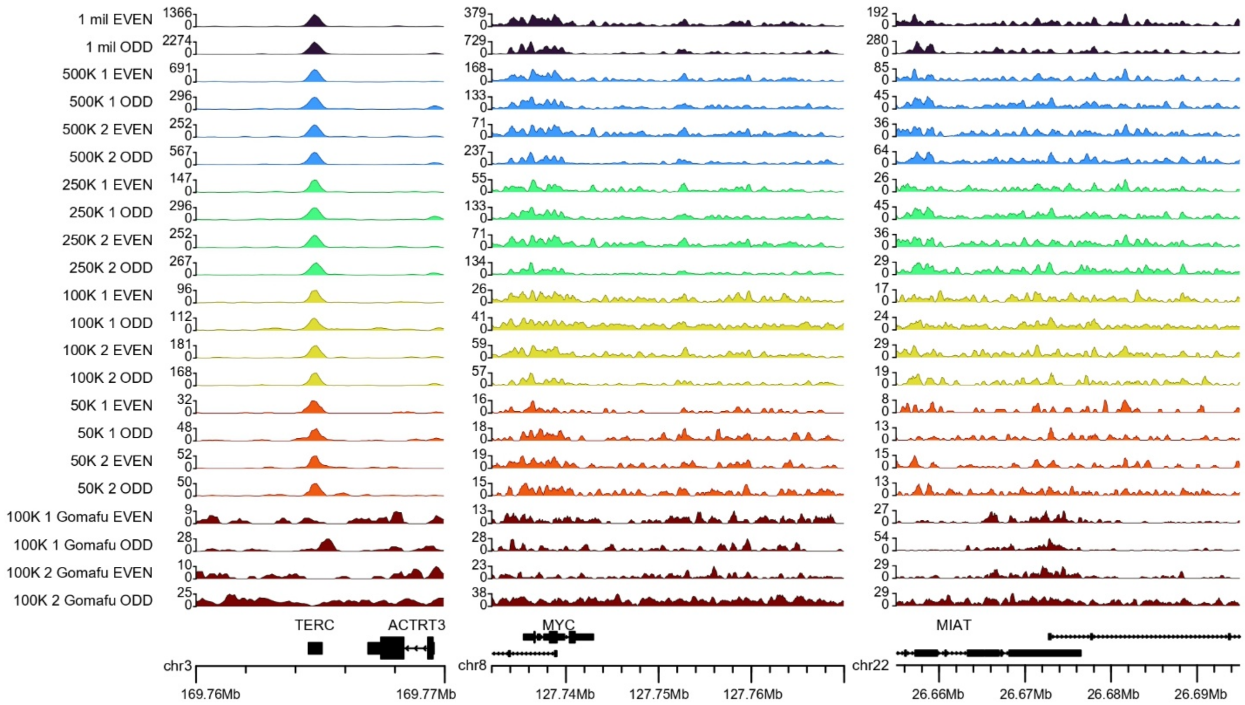


**Supplementary Figure 5. *TERC* and *GOMAFU* binding in HeLa cells by muChIRP-seq, even and odd reproducibility.** Even and odd genome browser tracks show *TERC* and *GOMAFU* binding in HeLa cells across replicates of decreasing input quantities.

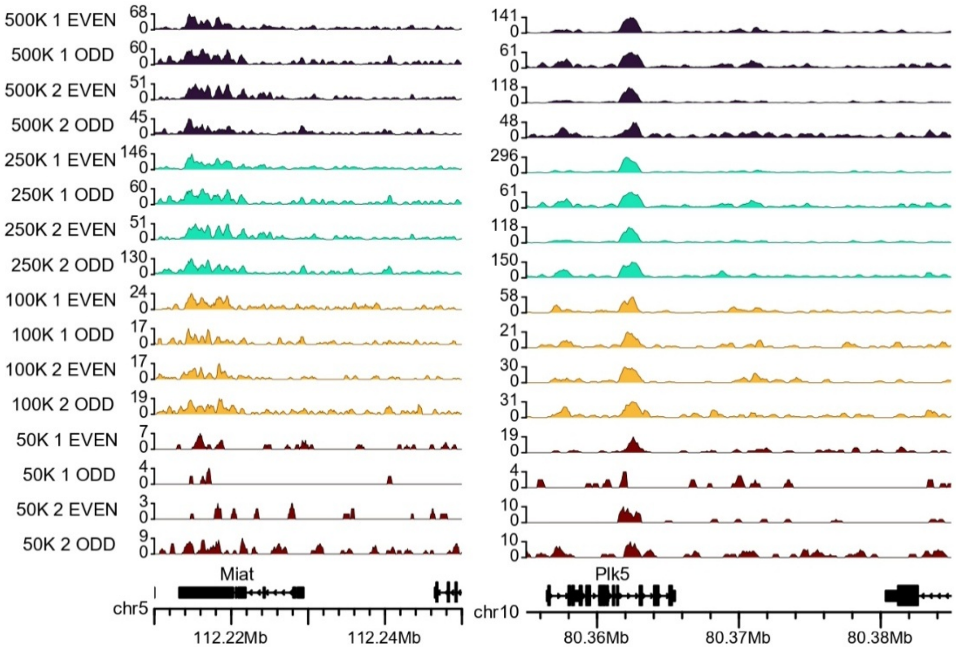


**Supplementary Figure 6. *Gomafu* binding in NE-4C cells by muChIRP-seq, even and odd reproducibility.** Even and odd genome browser tracks show *Gomafu* binding in NE-4C cells across replicates of decreasing input quantities.

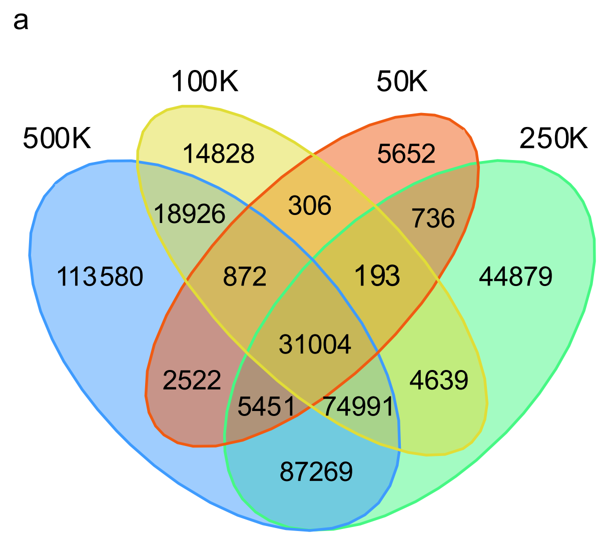


**Supplementary Figure 7. True peak overlap across input quantities.** Venn diagrams showing overlap of “true” peaks across input quantities (500K–50K). For each input level, peak sets represent the union of “true” peaks from both replicates for **(a)** *TERC* in HeLa and **(b)** *Gomafu* in NE-4C.

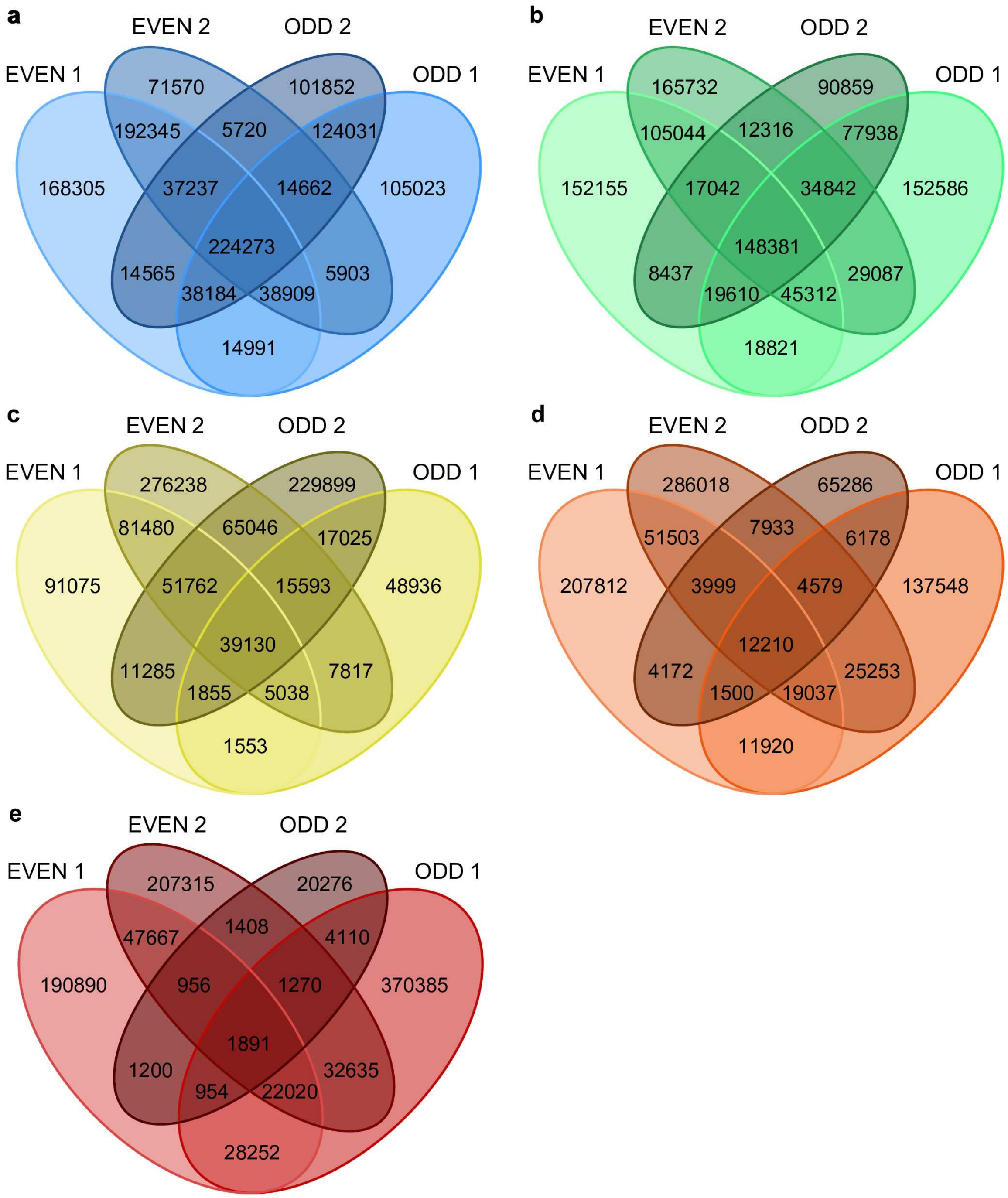


**Supplementary Figure 8. *TERC* and *GOMAFU* replicate peak overlap across input quantities for even and odd probe sets in HeLa cells.** Venn diagrams showing overlap of peaks between even and odd replicates across input quantities (500K–50K) for *TERC* and *GOMAFU* in HeLa: **(a)** 500K cells; *TERC* **(b)** 250K cells; *TERC* **(c)** 100K cells; *TERC* **(d)** 50K cells; *TERC* **(e)** 100K cells; *GOMAFU*.

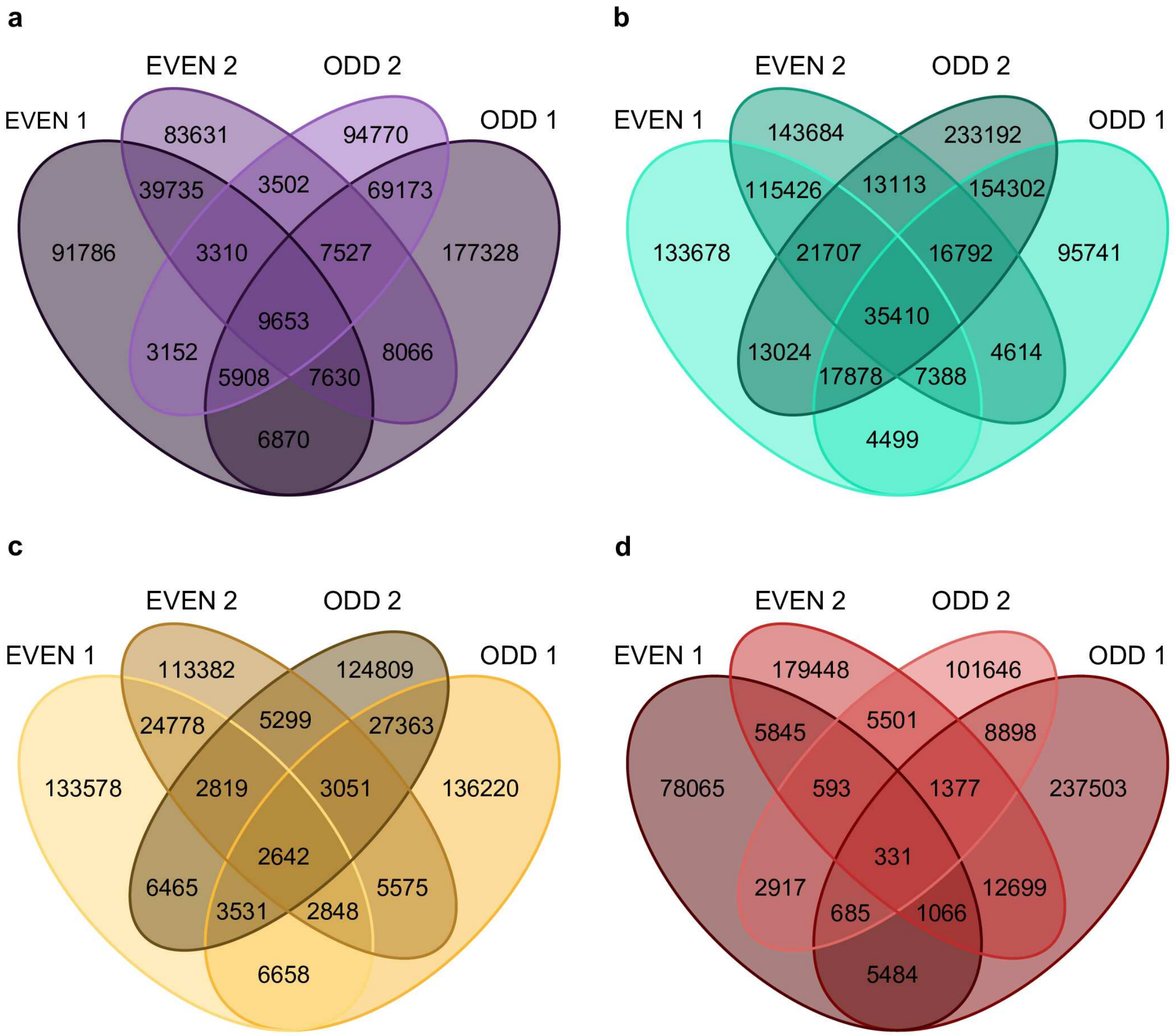


**Supplementary Figure 9. *Gomafu* replicate peak overlap across input quantities for even and odd probe sets in NE-4C cells.** Venn diagrams showing overlap of peaks between even and odd replicates across input quantities (500K–50K) for *Gomafu* in NE-4C: **(a)** 500K cells **(b)** 250K cells **(c)** 100K cells **(d)** 50K cells.

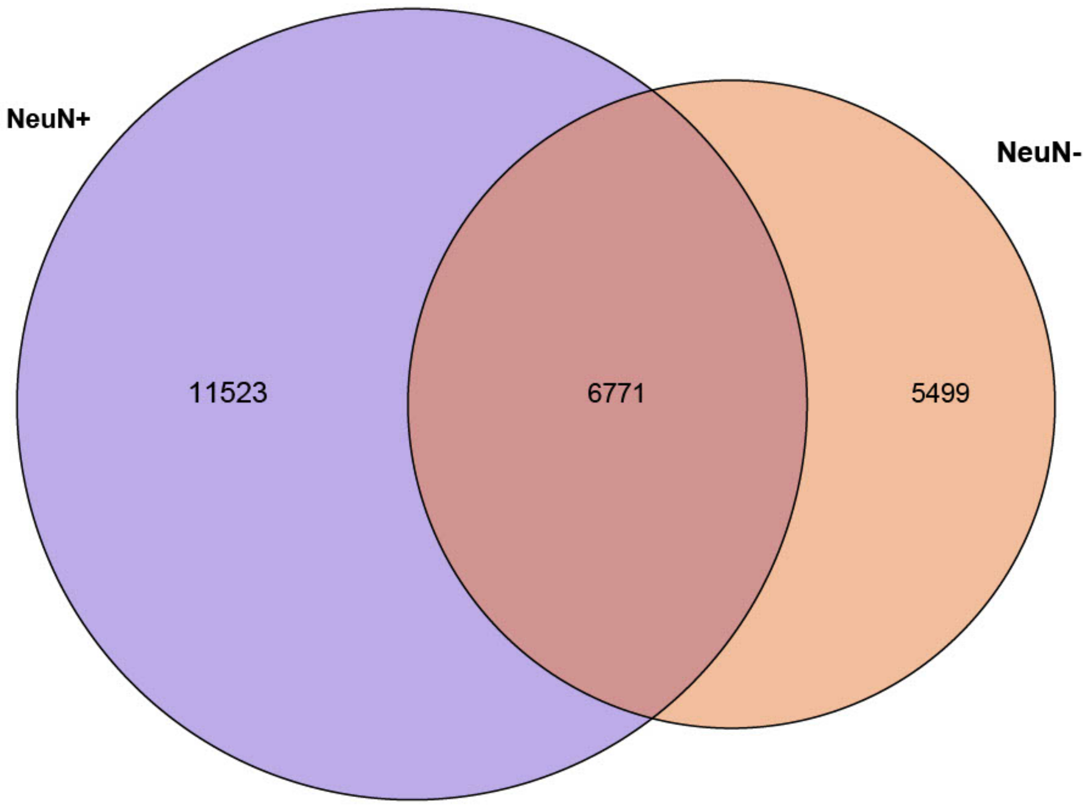


**Supplementary Figure 10. NeuN+ and NeuN- peak overlap in mouse cortex.** Venn diagram showing overlap of “true” peaks between NeuN+ and NeuN- samples. Peak sets for each cell type represent the overlapping “true” peaks between both replicates.

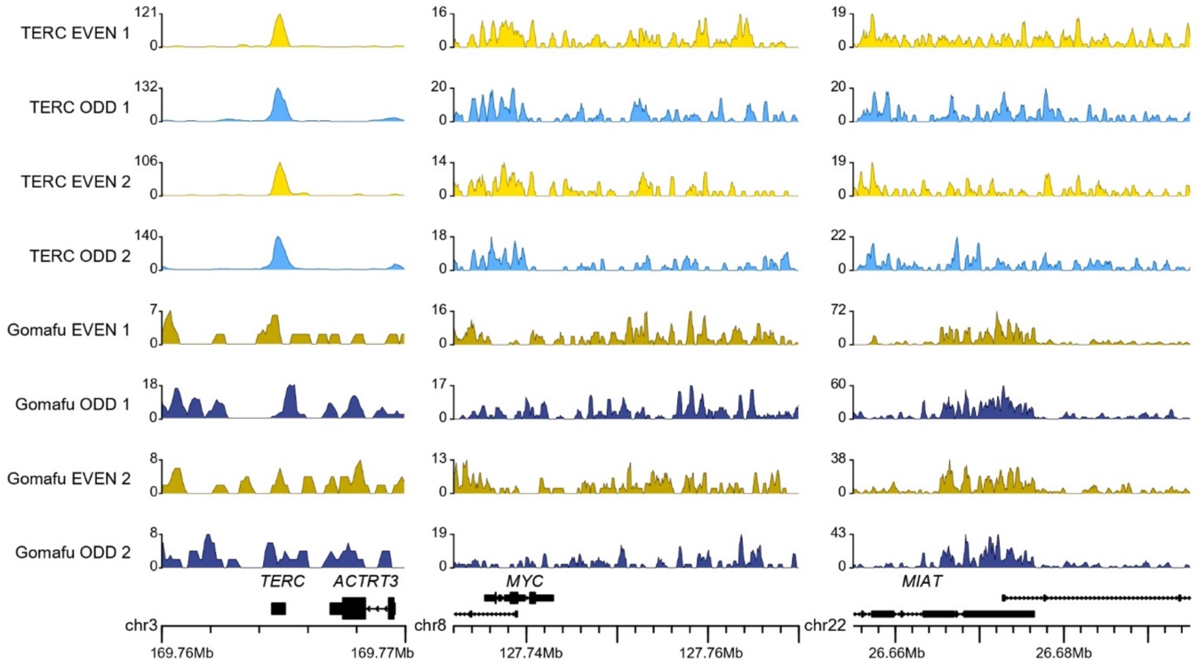


**Supplementary Figure 11. Genome browser tracks for *TERC* and *GOMAFU* muChIRP replicates in CNTRL-24.** Genome browser tracks showing replicate muChIRP even and odd probe pool libraries for TERC and *GOMAFU* in NeuN+ nuclei from postmortem brain sample CNTRL-24. Representative genomic regions are shown for each lncRNA.

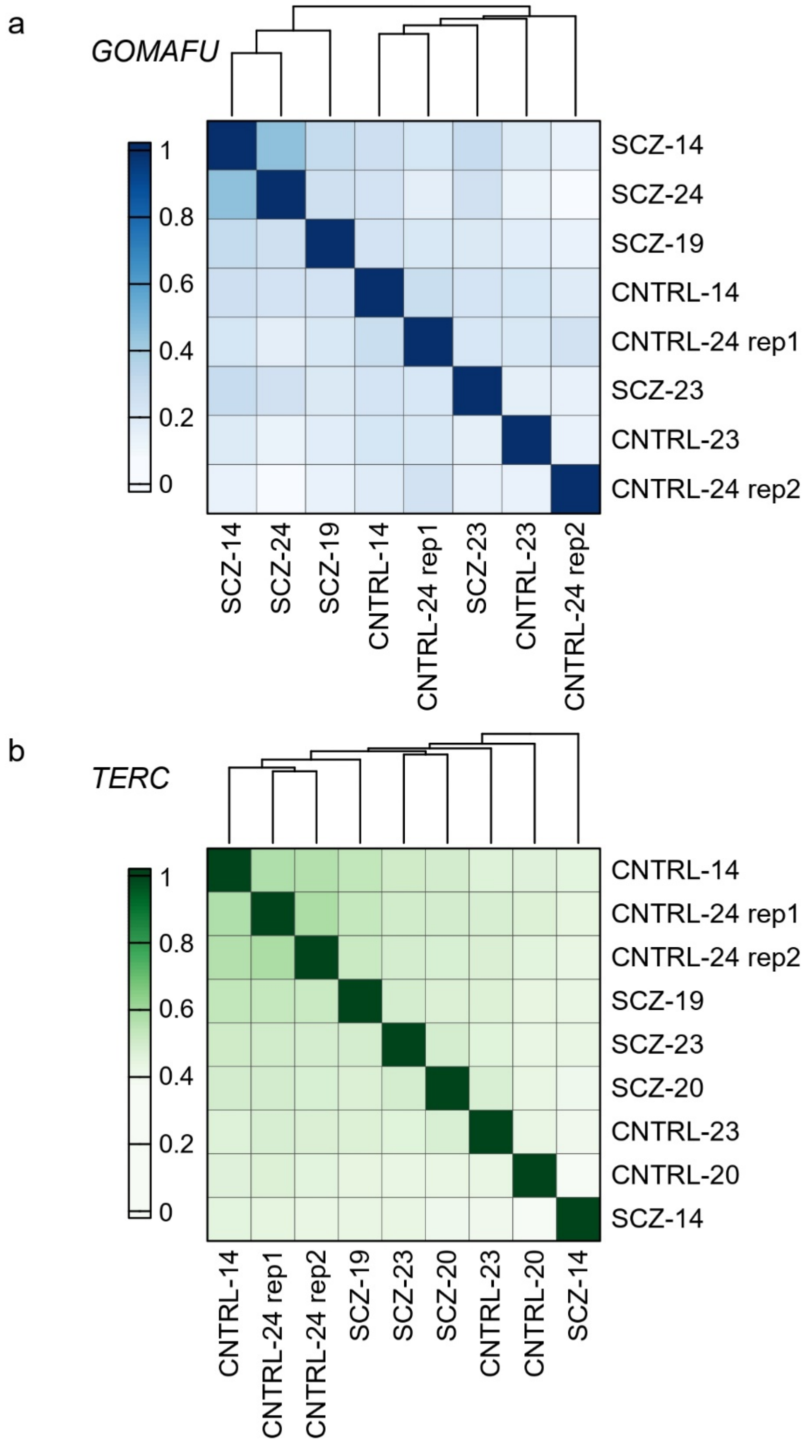


**Supplementary Figure 12. Pearson Correlation for *TERC* and *GOMAFU* in human prefrontal cortex.** Pearson correlation of signal over all true peak locations for (a) *GOMAFU* and (b) *TERC* samples. Hierarchical clustering determined order of samples displayed and is indicated by guidelines at the top of each heatmap.

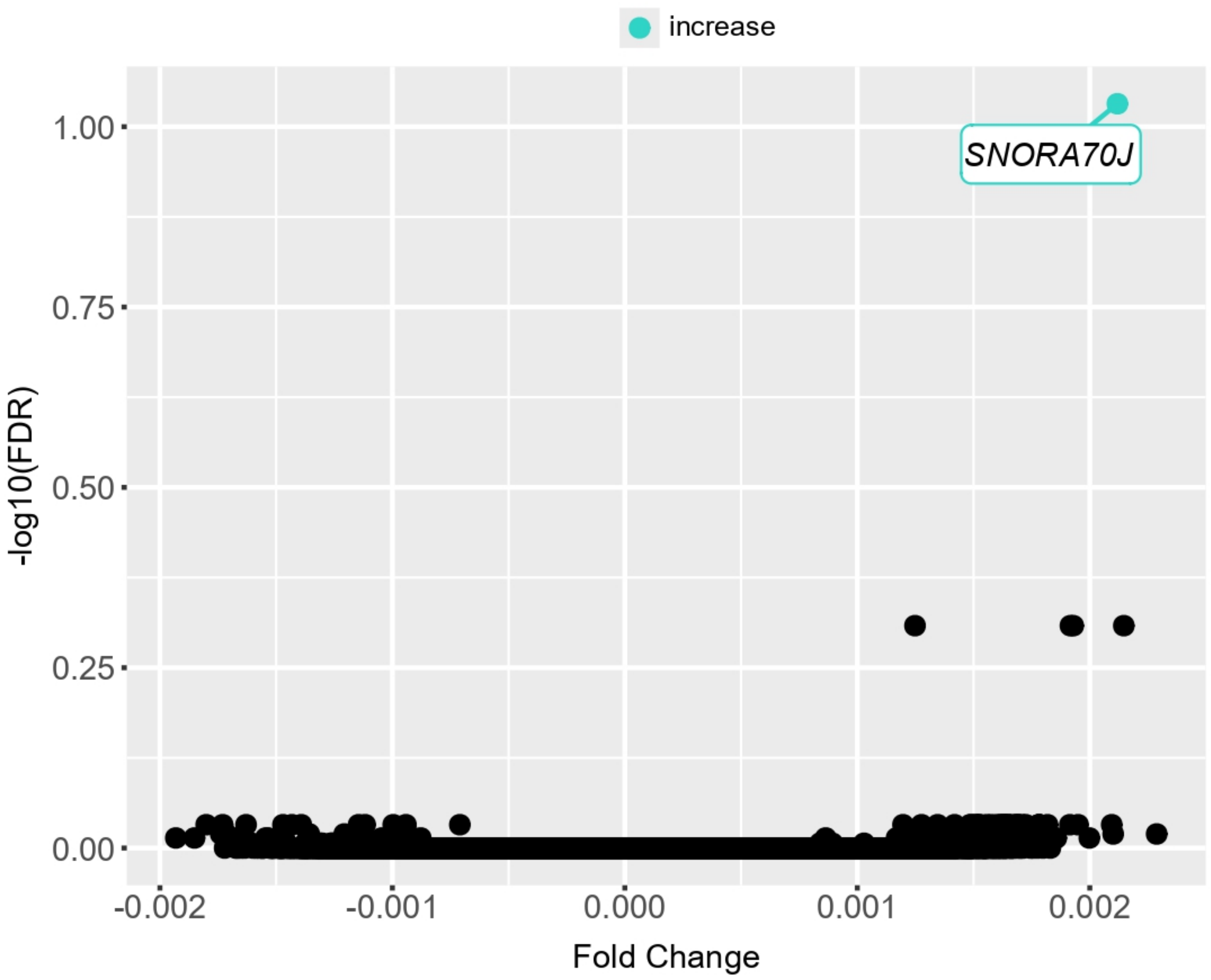


**Supplementary Figure 13. Volcano plot of differential *TERC* peaks in schizophrenia versus control.** Peaks are separated by increased or decreased enrichment in schizophrenia; there is only one significantly differential *TERC* peak (FDR ≤ 0.2) and it has increased enrichment in schizophrenia. This peak is labeled with the corresponding gene name.

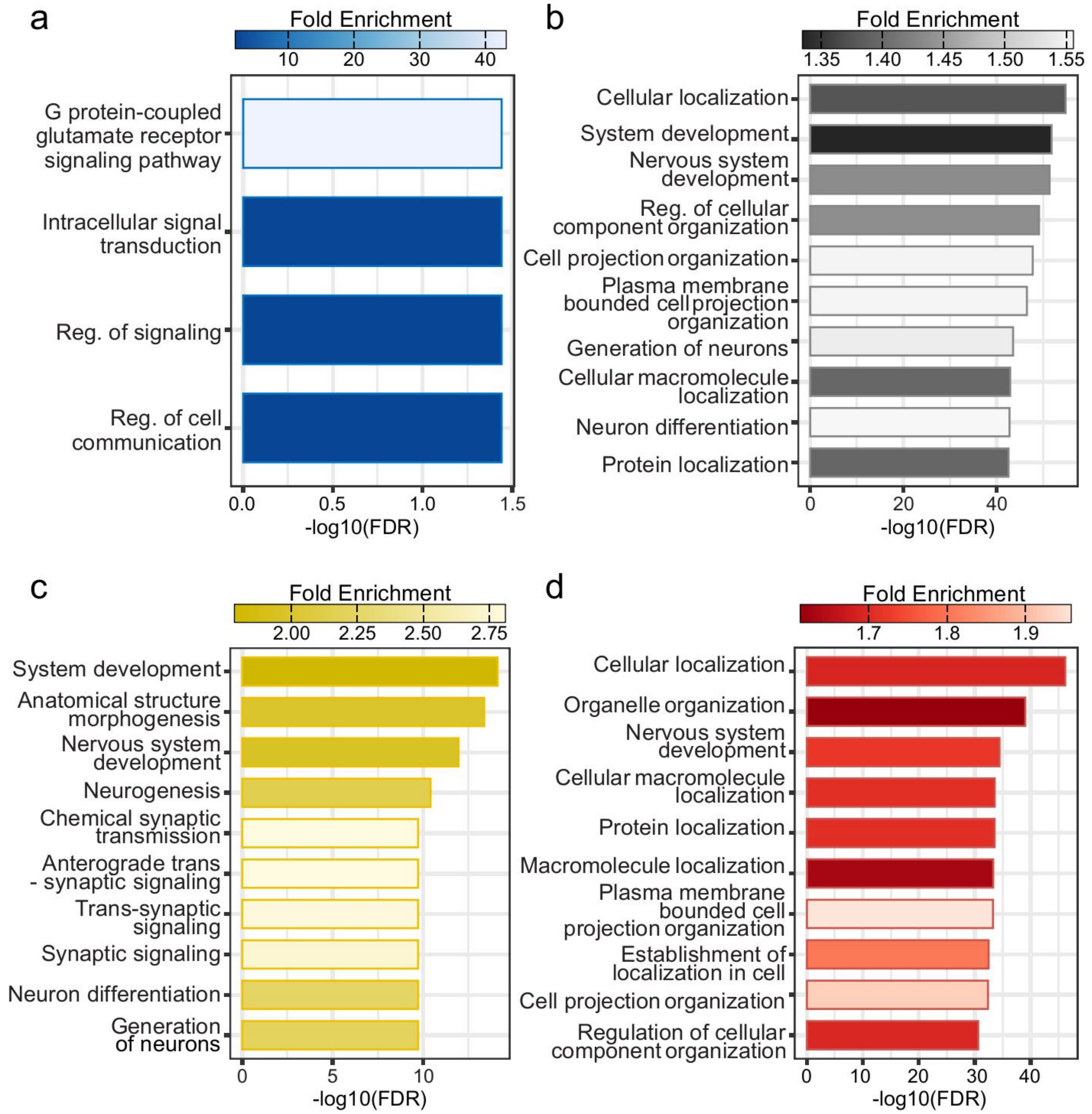


**Supplementary Figure 14. GO Biological Processes in differential multi-omic datasets.** Significantly enriched (FDR < 0.05) GO biological processes among differential peaks or genes for (a) *GOMAFU*, (b) H3K4me3, (c) H3K27ac, and (d) RNA-seq datasets. There were no significantly enriched GO biological processes for *TERC*.

**Supplementary Data**

**Supplementary Data 1. Postmortem human brain sample information.**

**Supplementary Data 2. Human *GOMAFU*, mouse *Gomafu*, and mouse *Terc* probe designs.**

**Supplementary Data 3. Metadata of all muChIRP-seq datasets.**

**Supplementary Data 4. Full list of differential peaks and genes identified on postmortem samples (FDR <0.2).**

**Supplementary Data 5. Full list of KEGG and GO terms identified by differential analysis and ShinyGO (FDR <0.05).**
